# Isoform-specific Patterns of Phosphorylation on Axonemal Dynein Heavy Chains

**DOI:** 10.1101/2025.03.10.642390

**Authors:** Miho Sakato-Antoku, Ramila S. Patel-King, Kazuo Inaba, Jeremy L. Balsbaugh, Stephen M. King

## Abstract

Axonemal dyneins power ciliary motility and phosphorylation of key intermediate and light chain components affects the regulation and properties of these motors in very distantly related organisms. It is also known that many axonemal dynein heavy chains are subject to this post-translational modification although this has been little studied. Here we examine axonemal dynein heavy chains from a broad range of ciliated eukaryotes and identify phosphorylated sites embedded within various kinase recognition motifs such as those for PKA, PKC and casein kinase II. Mapping these sites onto discrete heavy chain types reveals class-specific locations apparently mediated by different kinases. For example, we find that all *Chlamydomonas* α heavy chain phosphorylation sites are in an extended loop derived from AAA5 that arches over the coiled-coil buttress which in turn interacts with the microtubule-binding stalk. In contrast, most sites in the monomeric inner arm dyneins occur very close to the N-terminus and may be involved in assembly processes. In *Chlamydomonas*, the two cilia (termed *cis* and *trans*) exhibit different intrinsic beat frequencies and we identify cilium-specific phosphorylation patterns on both the α heavy chain and outer arm docking complex consistent with differential regulation of these motors in the two organelles.

**Significance:**

- Many axonemal dynein heavy chains are extensively phosphorylated. However, the location, sequence context and potential role of these modifications is uncertain.
- Here we use mass spectrometry to define phosphorylated sites in inner and outer arm dyneins from a broad range of motile ciliated eukaryotes.
- We identify numerous modified sites that correspond to various kinase consensus sequences and reveal that multiple sites occur in conserved locations depending on dynein heavy chain class. We also find phosphorylation modifications that are specific to either the *cis* or *trans* cilium of *Chlamydomonas* which may be related to their inherent beat frequency imbalance.

## Introduction

Motile cilia power the movement of individual cells and generate fluid flows over surfaces and within cavities. The cellular machinery underlying ciliary motion is the axoneme which generally consists of a ring of nine outer doublet microtubules with dynein motors arranged in two rows that generate inter-doublet sliding, a central pair of microtubule singlets, and radial spokes which transmit signals from the central pair apparatus to the dynein arms as well as numerous additional components (Walton et al., 2023). The beating of these organelles requires precise control of the linear arrays of dynein motors such that waves of activity propagate along the ciliary outer doublet microtubules and the region of active sliding switches so as to generate forward and reverse bends (Lin and Nicastro, 2018). In addition, cilia can alter their beat frequency and waveform in response to various signaling inputs such as changes in Ca^2+^, cyclic nucleotides and redox state (Habermacher and Sale, 1997; Hamasaki et al., 1991; Harrison et al., 2002; Kamiya and Witman, 1984; King and Dutcher, 1997; Wakabayashi and King, 2006). In humans, defective ciliary motility leads to the syndrome primary ciliary dyskinesia that involves multiple phenotypes (Reiter and Leroux, 2017; Shoemark and Harman, 2021) including infertility, severe respiratory problems, hydrocephalus (very rare), and *situs inversus*; ciliary motility defects are also associated with congenital heart disease (Li et al., 2015), various forms of epilepsy (Faubel et al., 2022; King, 2006) and several psychiatric disorders (Alhassen et al., 2021).

Axonemal dynein motors fall into three broad categories - the outer dynein arm which contains two (or in some organisms three) different HC motor units, the HC heterodimeric inner dynein arm I1/f and a series of monomeric HC motors that are classified into three distinct groupings (termed IAD-3, IAD-4 and IAD-5) (Wickstead, 2018). These motor proteins are tightly associated with a series of other intermediate and light chain components that vary depending on dynein type (King, 2018). The outer arms provide most of the power for ciliary beating while the inner arms control the precise waveforms and are important for beating under viscous load (discussed in (King et al., 2023)).

Post-translational modifications (PTMs) control the properties and dynamics of numerous cellular systems. Axonemal dyneins are subject to multiple post-translational alterations including N-terminal proteolytic processing and acetylation (Sakato-Antoku et al., 2023), methylation (Sakato-Antoku et al., 2024) and phosphorylation *e*.*g*. (Chilcote and Johnson, 1990; Habermacher and Sale, 1997; Hamasaki et al., 1991; King and Witman, 1994; Piperno and Luck, 1981); the latter two being potentially reversible modifications. These alterations might impact the cytoplasmic pre-assembly, axonemal incorporation and/or mechanochemical properties of individual motors. By analogy with tubulin, actin and histone PTM codes that modify functional properties (Janke and Magiera, 2020; Jenuwein and Allis, 2001; Mu et al., 2020), this has given rise to the concept of discrete post-translational dynein changes that might alter various properties of these highly complex enzymes (Sakato-Antoku et al., 2025).

In *Chlamydomonas*, phosphorylation of an intermediate chain (termed IC138) associated with the I1/f inner arm dynein affects the motile properties of cilia (Habermacher and Sale, 1997; King and Dutcher, 1997). When this IC is modified by cAMP-dependent protein kinase A (PKA) and/or casein kinase I (CK I), the I1/f dynein becomes inactive resulting in altered ciliary waveform but normal beat frequency. This effect is reversed by the action of two phosphatases (PP1 and PP2A) (Habermacher and Sale, 1996; Yang et al., 2000). However, it remains unclear how these modifications on an IC component actually impact the I1/f dynein HCs to alter motor activity. In *Paramecium*, the phosphorylation of an outer arm dynein light chain (termed p29) by PKA increases the rate of dynein-driven microtubule gliding *in vitro* (Barkalow et al., 1994). Furthermore, treating *Paramecium* with a membrane-permeable cAMP analogue led to the phosphorylation of the p29 light chain and increased cell swimming velocity (Hamasaki et al., 1995). Again though, the mechanism involved in controlling HC motor activity *via* this light chain modification remains uncertain. Furthermore, phosphorylation of a Tctex2 light chain in salmonid axonemal dyneins has been linked to the activation of sperm motility (Inaba et al., 1999). There is also evidence for the phosphorylation of axonemal dyneins directly on the HCs. *In vivo* long-term radio-labeling with ^32^P-phosphate followed by biochemical fractionation of isolated cilia (King and Witman, 1994; Piperno and Luck, 1981) as well as multiple whole cilia mass spectrometry studies (Pan et al., 2011; Wagner et al., 2006; Wang et al., 2014) in *Chlamydomonas* identified several phosphorylated dynein HCs; pulse-labeling of intact *Chlamydomonas* cells with ^32^P-phosphate suggested that some of these phosphorylated sites turn over rapidly within cilia *in vivo* (King and Witman, 1994).

Recently, we examined electrophoretically purified dynein HCs derived from members of four taxonomic supergroups (Alveolata, Chloroplastida, Discoba and Opisthokonta; see (Burki et al., 2020) for a recent eukaryotic tree-of-life illustrating their relationships) to identify conserved sites of methylation using tandem mass spectrometry (Sakato-Antoku et al., 2025). Here we have searched for phosphorylated residues in dynein HCs from this broad range of motile ciliated eukaryotes. We find that many dynein HCs are phosphorylated at key sites involved in assembly and mechanochemistry and that the specific sites modified vary considerably depending on the axonemal dynein HC class. Furthermore, motif analysis suggests that multiple different kinases controlled by various signaling inputs such as Ca^2+^/calmodulin and cAMP are required for these modifications, leading to the concept that multiple regulatory inputs may be integrated at the level of individual dynein motors.

## Results

### Phosphorylation of dynein heavy chains in motile ciliated eukaryotes

Dynein HCs were electrophoretically isolated from cilia or other axonemal dynein-containing samples from *Ceratopteris richardii* (water fern, tracheophyte), *Chlamydomonas reinhardtii* (green alga, chlorophyte), *Ciona intestinalis* (sea squirt, chordate), *Crassostrea gigas* (Pacific oyster, mollusk), *Drosophila melanogaster* (fruit fly, dipteran insect), *Drosophila willistoni* (fruit fly, dipteran insect), *Hemicentrotus pulcherrimus* (sea urchin, echinoderm), *Mnemiopsis leidyi* (comb jelly, ctenophore), *Oncorhynchus mykiss* (rainbow trout, actinopterygian), *Rattus norvegicus* (Norway rat, mammal), *Takifugu rubripes* (puffer, actinopterygian), *Tetrahymena thermophila* (ciliate, alveolate) and *Trypanosoma brucei* (trypanosome, kinetoplastid) as described in (Sakato-Antoku et al., 2025). The HCs were then digested with trypsin and/or endoproteinase Asp-N and subjected to high-resolution tandem mass spectrometry for peptide identification using pSer, pThr and pTyr modifications. This analysis (Table 1 and see Table S1 for full details of mass spectral coverage and all modified residues identified) found a total of 82 phosphorylated sites (66 pSer and 16 pThr) in 49 of the 79 identified axonemal HCs from *Chlamydomonas, Ciona, Crassostrea, Hemicentrotus, Oncorhynchus, Rattus, Takifugu, Tetrahymena* and *Trypanosoma*; it also identified a single pSer residue on the rat cytoplasmic dynein HC. Furthermore, previous mass spectrometric analyses of various *Chlamydomonas* cilia samples by other groups have found 22 pSer and 5 pThr HC residues in addition to the ones identified by our study (see data from (Pan et al., 2011; Wagner et al., 2006; Wang et al., 2014) that are included in Tables 1 and S1). In contrast, no evidence for phosphorylated sites was obtained for a combined 31 axonemal dynein HCs from the fern, insect and ctenophore samples.

**Table 1.** Phosphorylation Sites Identified in Axonemal Dynein Heavy Chains from Motile Ciliated Eukaryotes.

| Organism | Outer Arm |  |  | Inner Arm I1/f |  | Monomeric Inner Arms |  |  |  |
| --- | --- | --- | --- | --- | --- | --- | --- | --- | --- |
| | $\alpha$ HC* | $\beta$ HC | $\gamma$ HC | 1α HC | 1β HC | IAD-3 | IAD-4 | IAD-5 | Uncertain<sup>#</sup> |
| *Ceratopteris richardii* | --- | --- | --- | 0 | 0 | 0 | 0 | 0 | --- |
| *Chlamydomonas reinhardtii* | 6 | 1 | 0 | 1 | 0 | 18 | 4 | 6 | 12 |
| *Ciona intestinalis* | --- | 0 | 2 | 0 | 0 | 0 | 0 | 0 | --- |
| *Crassostrea gigas* | --- | 1 | 2 | 1 | 0 | 4 | 2 | 2 | --- |
| *Drosophila melanogaster* | --- | 0 | 0 | 0 | 0 | 0 | 0 | 0 | --- |
| *Drosophila willistoni* | --- | 0 | 0 | 0 | 0 | 0 | 0 | 0 | --- |
| *Hemicentrotus pulcherrimus* | --- | 0 | 2 | 0 | 1 | 4 | 0 | 3 | --- |
| *Mnemiopsis leidyi* | --- | 0 | 0 | 0 | 0 | 0 | 0 | 0 | --- |
| *Oncorhynchus mykiss* | --- | 0 | 0 | 0 | 0 | 5 | 1 | 0 | --- |
| *Rattus norvegicus* | --- | 2 | 0 | 0 | 0 | 1 | 2 | 0 | --- |
| *Takifugu rubripes* | --- | 0 | 0 | 0 | 1 | 3 | 0 | 0 | --- |
| *Tetrahymena thermophila* | 1 | 0 | 0 | 0 | 1 | 1 | 0 | 0 | --- |
| *Trypanosoma brucei* | --- | 4 | 0 | 0 | 0 | 1 | 2 | 4 | --- |

\* This outer arm HC is present in *Chlamydomonas* and *Tetrahymena* (where it is termed the $\gamma$ HC); metazoans, ferns and trypanosomes do not encode an ortholog.

### Dynein heavy chains are phosphorylated on serine and threonine residues

Mass spectral analyses of dynein HC-derived peptides provide evidence for a combined 109 pSer/pThr phosphorylation sites on various axonemal dynein HCs (Fig. 1). Two putative phosphotyrosine (pY)-containing peptides were identified in the informatics searches for inner arm dyneins *Oncorhynchus* DNAH2 (CARmePVLpYR_me2_K) and *Tetrahymena* TTHERM_001151438 (LAQTLYIDpYEPLNR). However, following manual validation through visual inspection, these were determined to be very low confidence or false positive identifications. Thus, the current data support the phosphorylation of axonemal dynein HCs exclusively on Ser and Thr residues, with the majority of modifications occurring on Ser residues and only a few altered Thr residues in most dynein HC classes. This is consistent with thin-layer chromatography phospho-amino acid analysis of axonemal dynein polypeptides purified from *Chlamydomonas* cells labeled *in vivo* with [^32^P]-phosphate which identified mainly pSer with barely detectable amounts of pThr (King and Witman, 1994).

**Figure 1.**
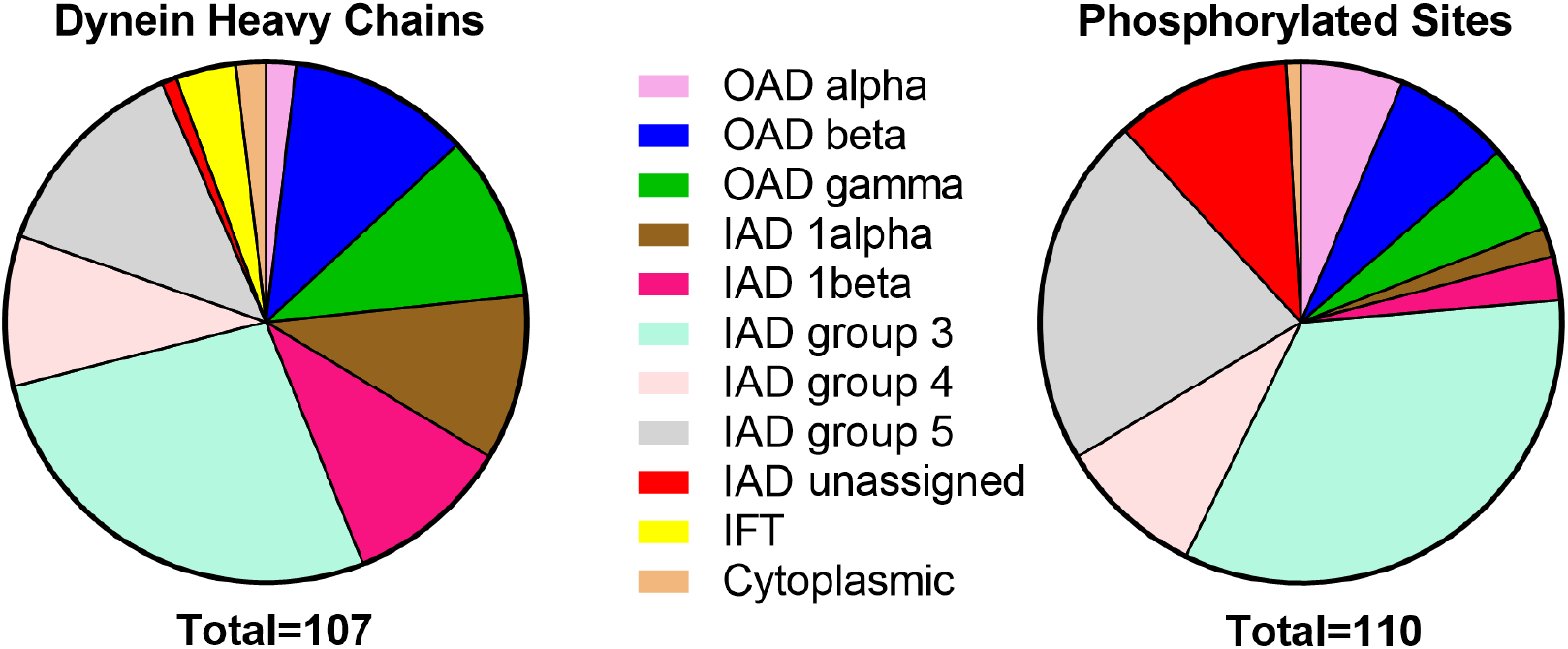
Distribution of Dynein Heavy Chain Classes and Phosphorylation Sites. The *left* sector plot displays the various dynein HC classes identified by mass spectrometry in samples derived from twelve motile ciliated eukaryotes (see Table 1; only one *Drosophila* species is included). The *right* sector plot illustrates the distribution of phosphorylated residues found in these various HC classes. This reveals that it is the monomeric inner arm HC groups that are most heavily modified, accounting for ∼75% of all sites identified.

### Phosphorylation of non-heavy chain dynein components

For *Chlamydomonas*, we were also able to search for phosphorylation of core non-HC dynein components in whole cilia samples. No phosphorylation modifications were identified on outer arm intermediate or light chains or non-HC components of the single motor inner arm dyneins which is consistent with *in vivo* ^32^P-phosphate labeling experiments (King and Witman, 1994; Piperno and Luck, 1981). However, we did find multiple pSer and pThr phosphorylation sites on the outer arm docking complex components DC1 (S_33_, T_73_, T_351_, T_498_, T_508_, S_510_, S_512_, S_514_, S_521_, S_628_, T_698_, S_700_ and S_709_) and DC2 (S_12_, S_15_, T_20_, S_266_, S_267_, T_512_ and S_548_); DC3 was not phosphorylated. The extensive phosphorylation of DC1 may be partly reflected in its abnormal migration on SDS-gels that was observed by (Takada and Kamiya, 1994). For inner arm I1/f components, we found a single pThr (T_682_) on the IC138 intermediate chain, a known regulatory phospho-protein, and also a single pSer (S_19_) on the ankyrin repeat protein FAP120 (DII7) that associates with IC138.

### Kinase motif analysis suggests multiple regulatory inputs

Several protein kinases, including cAMP-dependent protein kinase (PKA) and casein kinase I (CK I), have been linked to the phospho-regulation of the IC138 component of inner arm dynein I1/f from *Chlamydomonas* and the control of microtubule sliding rates (Habermacher and Sale, 1997; King and Dutcher, 1997; Yang and Sale, 2000); the phosphorylated site (T_682_) we identified in IC138 is embedded in the sequence _676_AAARPV**<u>pT</u>**AESAAA_688_ which appears more similar to a Ca^2+^/calmodulin-dependent protein kinase (CaMK II) motif than to PKA or CK I. PKA has also been found to modify the p29 light chain of *Paramecium* outer arm dynein and regulate its activity in a cAMP-dependent manner (Hamasaki et al., 1991). To assess the potential kinases involved in modifying axonemal dynein HCs, we analyzed the sequences surrounding identified phosphorylated residues for known kinase recognition motifs (Fig. 2 and Table S2). This revealed a clear division between outer and inner arm dyneins and especially between the outer arms and the single HC inner arm motors.

**Figure 2.**
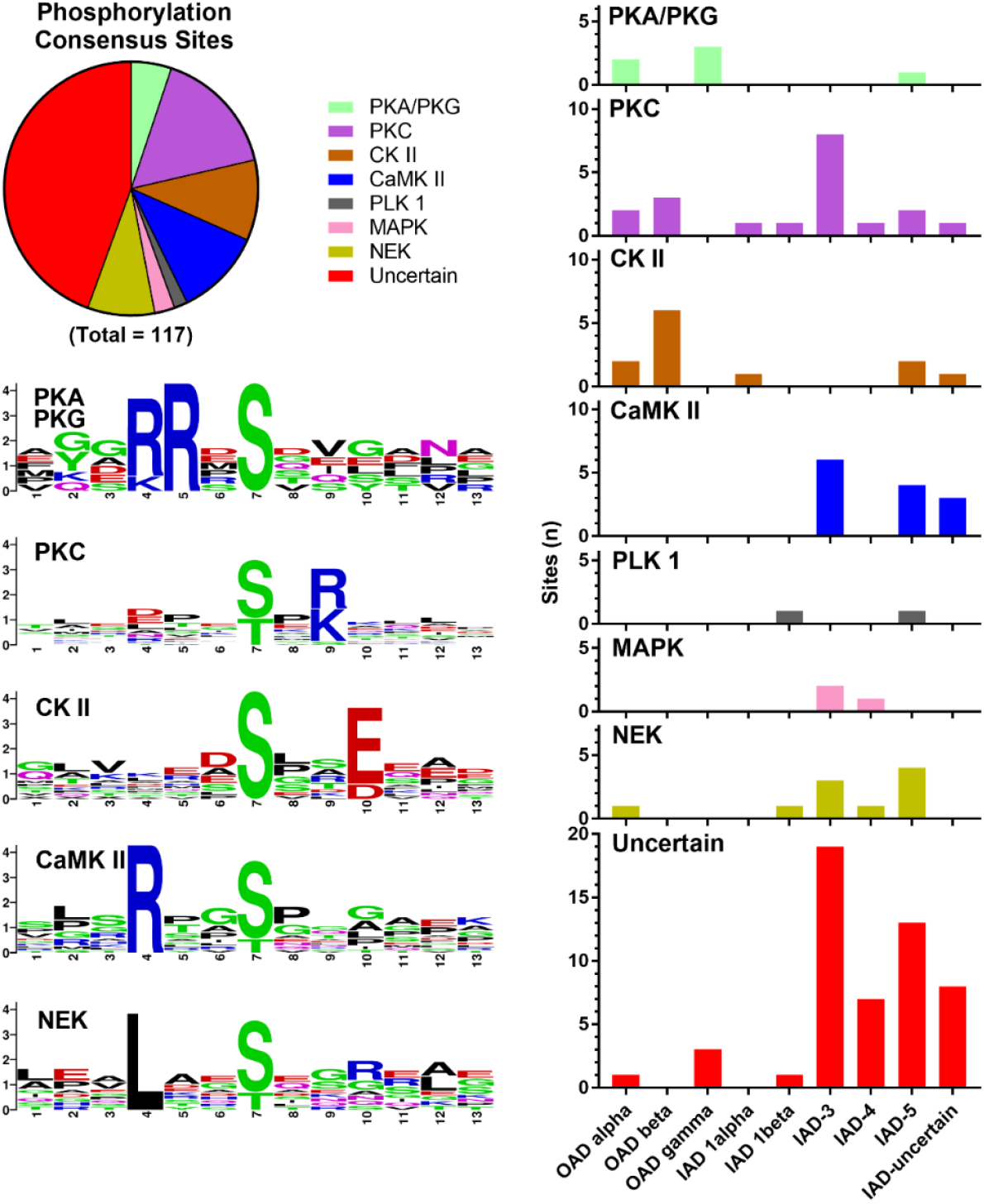
Kinase Motif Analysis of Dynein Heavy Chain Phosphorylation Sites. The sector plot (*upper left*) illustrates the distribution of the various kinase motifs identified; their occurrence in the different dynein classes is shown at *right*. Approximately half of the phosphorylated sites do not conform to any of the standard motifs; nearly all of these were found in the monomeric inner arms. In a few cases, a sequence conformed to more than one consensus motif. WebLogo outputs for five of the identified motifs are shown at *lower left*. In combination, this analysis reveals a stark division between the multimeric and monomeric dynein HC motors. The individual sequence motifs in which the identified phosphorylated residues are embedded are provided in Table S2 arranged by HC class.

Outer arm phosphorylated sites mostly conformed to the consensus sequences for PKA, protein kinase C (PKC) and casein kinase II (CK II); the PKC and CK II sites were identified in the paralogous α and β HCs, while PKA sites and those of uncertain origin were present in γ HC orthologs illustrating the potential for differential control of each motor within individual dynein arms. In contrast, the inner arm sites were found to represent PKC, CaMK II, and NimA-related kinase (NEK) motifs with a few examples of Polo-like kinase 1 (PLK-1), MAP kinase (MAPK) and PKA motifs. However, the majority of monomeric inner arm HC sites did not conform to any of these consensus sequences and consequently, the kinase(s) likely responsible for these modifications are uncertain. Intriguingly, use of the Kinase Predictor v.0.8 tool which employs sequence matrices based on a large-scale analysis of human kinase substrates (Sugiyama et al., 2019), revealed that several of these uncertain inner arm motifs which include the sequence [S/T]P may be recognized by one or more cyclin-dependent kinases (CDKs).

Analysis of phosphorylated docking complex components using the Kinase Predictor also suggests multiple kinases act on DC1 and/or DC2, including CK II, PKC, CaMK I and at least one CDK. Although the inner arm protein IC138 is reported to be phosphorylated by CK I (Yang and Sale, 2000), we did not identify any phosphorylated HC sites conforming to the CK I consensus sequence [D/E][D/E][D/E]xx[S/T]ϕ or to the pre-phosphorylated motif [pS/pT]xx[S/T]ϕ that is also a target for this enzyme.

Together, these observations imply that dynein phosphorylation status likely responds to numerous distinct kinases that impact on discrete HC classes and/or other components and provides a potential mechanism for signal integration at the level of individual motor units within a given dynein arm.

### Dynein heavy chain class-specific phosphorylated regions

Mapping the locations of all the identified phosphorylated sites onto maps of HC domain organization clearly reveals some striking differences between the various HC isoform groups (Fig. 3). For example, all the phosphorylated residues found in the outer arm α HC are located in a loop region that extends from AAA5; the paralogous β HC lacks this extended loop segment. Clusters of sites are present in similar loop locations in inner arm dynein group 5 HCs. In contrast, all HCs except the outer arm α HCs have one or more phosphorylation sites near the N-terminus, and indeed for the monomeric inner arm groups the majority (22/37) of sites identified are located within ∼200 residues of the N-terminus. The outer arm β and γ HCs and inner arm I1/f 1α HC also have one of more phosphorylation sites positioned between the HC-HC interaction sites (DHC-1N domains) and the linker regions; for the β and 1α HCs these are PKC or CK II motifs while for the γ HC they are PKA motifs or the product of an uncertain kinase. This analysis also revealed that shared phosphorylation motifs exhibit a restricted occurrence. For example, with only two exceptions, all the CaMK II sites are located in the regions N-terminal of the linker domains. Similarly, all identified PKA sites occur either in the N-terminal regions or in AAA5.

**Figure 3.**
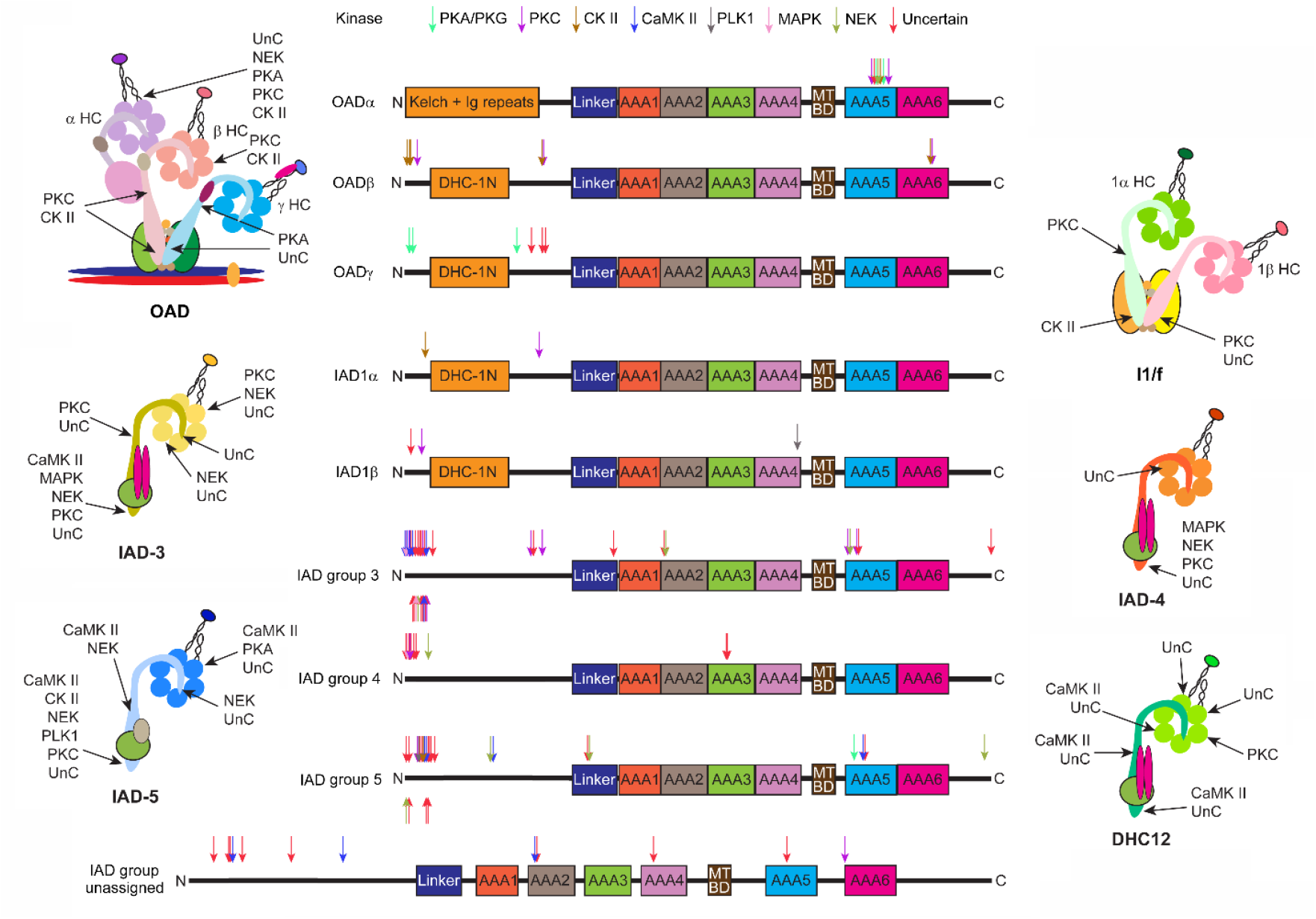
Location of Axonemal Dynein Heavy Chain Phosphorylation Sites. The location of all phosphorylated sites found in various dynein HC classes from all organisms examined are shown on the generic domain organization maps which indicate the AAA domains, microtubule-binding domain (MTBD) and the N-terminal regions involved in HC-HC associations - “DHC-1N” in outer arm β and γ HCs and inner arm I1/f 1α and 1β HCs or “Kelch + Ig repeats” in the α HC. As different members of these classes have somewhat variable domain spacings the locations indicated are approximate. The arrows indicate the sites of phosphorylation identified in all analyzed members of each HC class and the colors code for the different kinase consensus motifs to which each site conforms. To the sides of the linear maps are diagrams of the dyneins to which each HC belongs illustrating the approximate locations of the phosphorylation sites identified within the assembled holoenzymes. Sites modified by uncertain kinases are indicated by UnC. The unassigned monomeric inner arm dynein (termed DHC12) is present only in *Chlamydomonas*.

### Inner arm heavy chain phosphorylation sites in unstructured N-terminal segments

Using AlphaFold 3 we next examined the structural properties of phosphorylated regions in the various HC classes focusing initially on the large clusters of sites identified in the monomeric inner arm HC groups. In most cases, these modifications occur in loop regions or segments with little or no predicted tertiary structure when the HC is modeled alone (Fig. 4 *upper row*). However, nearly all these sites are located within the N-terminal ∼200 residues of these HCs and thus are in locations that have the potential to interact with other inner arm dynein components including actin, centrin and orthologs of the dimeric light chain termed p28 in *Chlamydomonas* or DNALI1 in mammals (Braschi et al., 2022). We then modeled the N-terminal regions from members of each monomeric HC group in combination with an actin monomer and either the dimeric p28/DNALI1 light chain or centrin (depending on group) (Fig. 4 *lower row*). This revealed a marked difference between monomeric inner arm HCs. For *Chlamydomonas* DHC7 (IAD group 5), the modified HC region is predicted to wrap around the actin monomer generating two clusters of phosphorylated sites. One cluster is located at an interface between the HC and actin while the second is at the opposite actin face and appears to involve HC interactions with both actin and centrin. In contrast, the phosphorylated regions of *Oncorhynchus* DNAH7 (IAD group3) and *Crassostrea* DNAH1 (IAD group 4) with actin and DNALI1 occur in unstructured segments that apparently are not involved in intra-dynein interactions and thus presumably associate with some other axonemal component(s).

**Figure 4.**
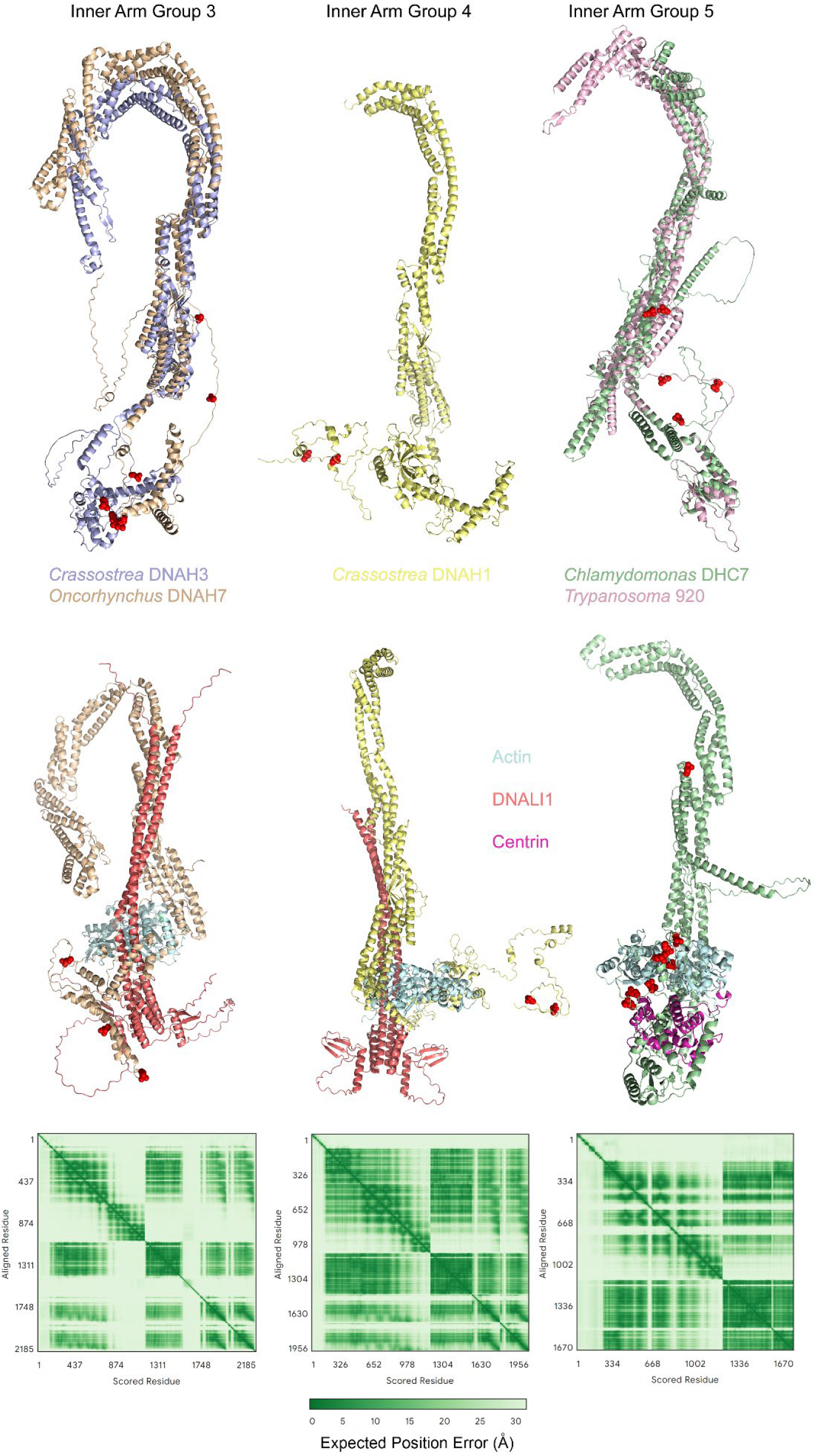
Phosphorylation of Monomeric Inner Arm Dynein Heavy Chain N-terminal Regions. The *upper row* shows AlphaFold3 models of the N-terminal 1050 residues from five phosphorylated monomeric inner arm HCs revealing the location of the modified residues in red (space-filling): inner arm group 3 HCs, *Crassostrea* DNAH3 (light blue), and *Oncorhynchus* DNAH7 (wheat); inner arm group 4 HC, *Crassostrea* DNAH1 (light yellow); and inner arm group 5 HCs, *Chlamydomonas* DHC7 (pale green) and *Trypanosoma* 920 (light pink). The structural models are oriented with the N-termini towards the bottom. Nearly all of the identified phosphorylated sites occur in loop regions. In the *lower row*, models for *Oncorhynchus* DNAH7 with actin and a DNALI1 dimer, *Crassostrea* DNAH1 with actin and a DNALI1 dimer, and *Chlamydomonas* DHC7 with actin and centrin are shown. For CrDHC7 in inner arm group 5 the phosphorylated HC regions now cluster around the actin monomer whereas in the other groups they do not. The AlphaFold3-generated plots at *bottom* show the expected position errors (in Å) for the three multimer models revealing that the locations of HCs, actin, and centrin or DNALI1 with respect to each other are well supported (lower predicted errors are indicated by darker green coloration). Sequences for *Chlamydomonas* proteins (DHC7, Cre14,g627576; actin; Cre13.g603700 and centrin, Cre11.468450) were downloaded from Phytozome, while those for *Oncorhynchus* DNAH7 (Q8WXX0), β-actin (A0A8C7RVS9) and DNALI1 (A0A8C7VEL0), and *Crassostrea* DNAH1 (A0A8W8J5H8), actin (017320), and DNALI1 (XP011434996) were from UniProt.

### Outer arm α and inner arm group 5 heavy chains contain an extended phosphorylated loop arching over the buttress

Studies in *Chlamydomonas* have revealed the outer arm α HC increases beat frequency and is implicated in the intrinsic beat frequency imbalance exhibited by the *cis* and *trans* cilia (*i*.*e*. those derived from the daughter and mother basal body, respectively) (Sakakibara and Kamiya, 1989; Sakakibara et al., 1991). The outer arm α HC orthologs from *Chlamydomonas* and *Tetrahymena* stand out by having a cluster of phosphorylated sites within the AAA5 domain; indeed, all identified sites on these isoforms occur in this one small region. In *Chlamydomonas*, the six identified α HC sites are within a segment of 45 residues that includes this loop; a single site was found in the equivalent region of the *Tetrahymena* ortholog (Table S1). Examination of the cryogenic electron microscopy structure of the *Tetrahymena* outer dynein arm (PDB 7K5B (Rao et al., 2021)) revealed that this modified region is missing from the structure presumably due to its inherent flexibility and thus poor electron density.

Analysis of this AAA4+AAA5 region of the *Chlamydomonas* α HC and the equivalent region of the paralogous β HC using AlphaFold 3 predicts an extended loop region in the α HC that is greatly extended compared to the short loop present in the β HC (Fig. 5); β HC orthologs from metazoans also lack this large loop. This extended phosphorylated loop arches over the buttress coiled coil that derives from AAA5; in the cilium it would be pointed away from the doublet microtubules and towards the ciliary membrane. Modeling of the entire α HC AAA ring suggests that this phosphorylated loop adopts a hairpin structure that overlays the buttress with three phosphorylated residues oriented towards the coil of the buttress outbound from AAA5 which might affect structural transitions in this region (Fig. S1 *upper panel*). In *Chlamydomonas*, the outer arm α HC has also been implicated in binding the dynein regulatory factor Lis1, the ciliary levels of which are dynamically modulated in response to viscous loading and other conditions that impact ciliary beat parameters (Pedersen et al., 2007; Rompolas et al., 2012). Modeling predicts that this regulatory protein may associate with the AAA ring directly adjacent to the extended phosphorylated loop (Fig. S1 *lower panel*).

**Figure 5.**
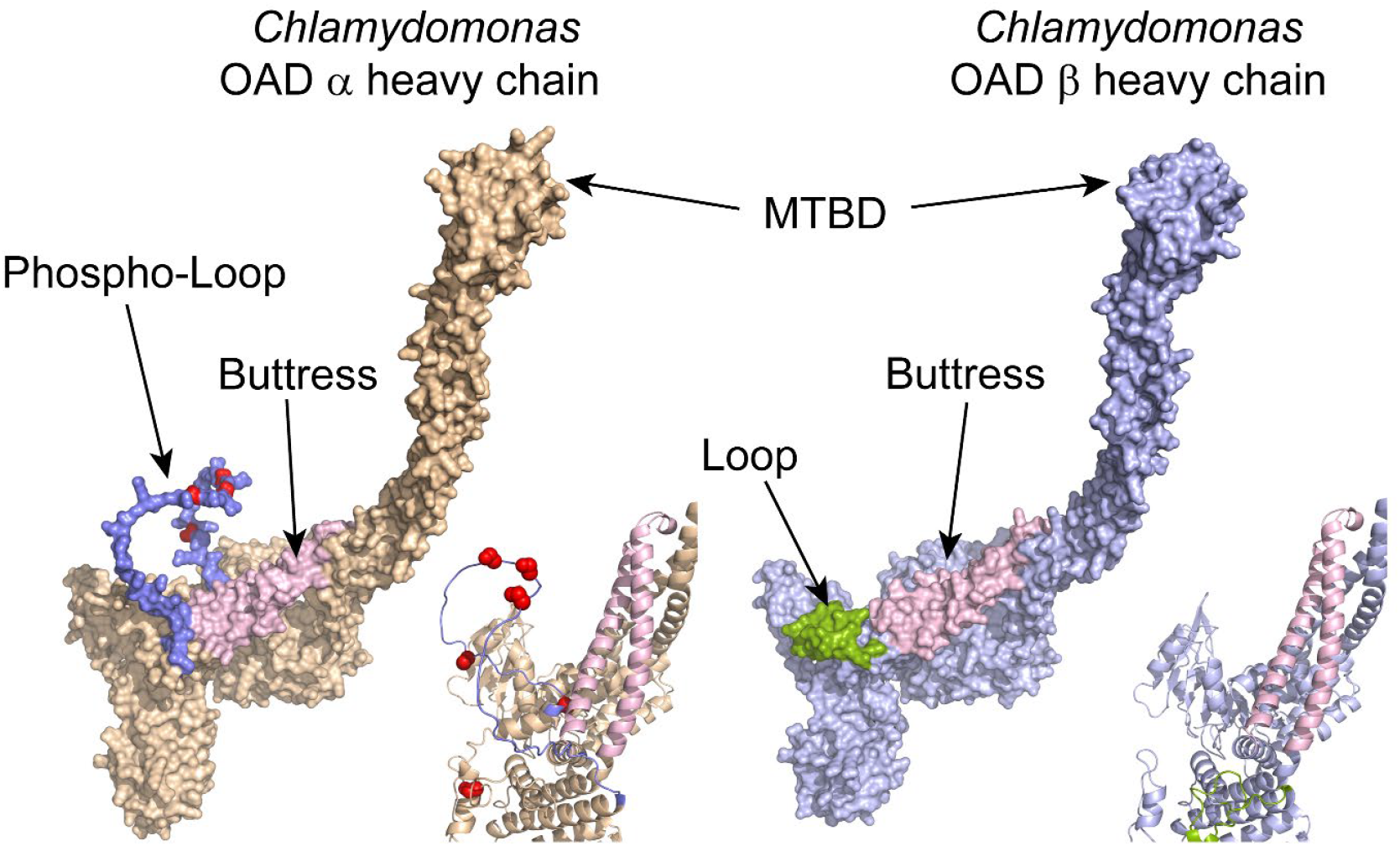
A Phosphorylated Loop Arches Over the Buttress from AAA5 in the Outer Arm α Heavy Chain. AlphaFold2 models (molecular surface and ribbon representations) for the *Chlamydomonas* outer arm dynein α and β HCs regions encompassing the microtubule binding and buttress segments derived from AAA4 and AAA5, respectively. In the α HC an extended disordered loop (slate blue) curves over the buttress (light pink) and is the site of five identified phosphorylated Ser residues (space-filling side chains in red). The equivalent much shorter loop in the β HC is shown in split-pea green.

Similarly, phosphorylated sites in inner arm group 5 HCs from *Trypanosoma* (11220) and *Chlamydomonas* (the DHC3 minor dynein HC) also occur in loops overarching the buttress although these loops are shorter than those in the *Chlamydomonas* α HC (Fig. 6 *left panel*); these inner arm HC loops point towards the central pair apparatus (see the arrangement of HCs in PDB 8J07 (Walton et al., 2023)). In contrast, *Chlamydomonas* DHC6, an inner arm group 3 HC, is phosphorylated at multiple sites directly on the buttress near their point of interaction with the coiled coil stalk from AAA4 that supports the microtubule-binding domain (Fig. 6 *right panel*).

**Figure 6.**
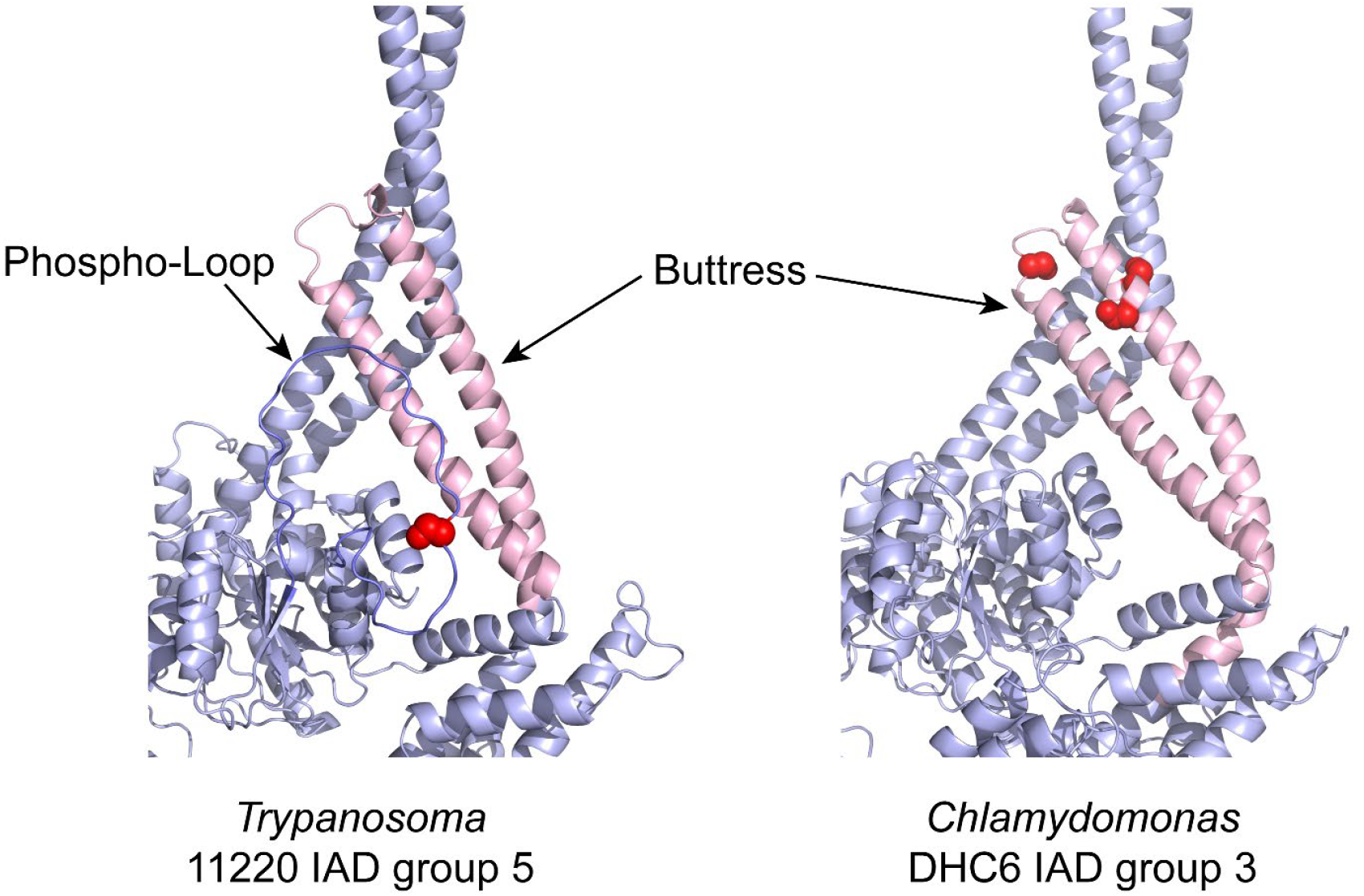
Loop and Buttress Phosphorylation on Monomeric Inner Arm Heavy Chains. AlphaFold 3 models of the buttress and phosphorylated loop from a *Trypanosoma* inner arm group 5 HC (11220), and a *Chlamydomonas* inner arm group 3 HC (DHC6) are shown. Similar to the outer arm α HC, the IAD group 5 HC has an extended phosphorylated loop that arches over the buttress. In contrast, the IAD group 3 HC is phosphorylated directly on buttress residues near the interaction site with the antiparallel coiled coil that supports the microtubule-binding domain. The buttress structure is colored light pink, the phosphorylated loop in slate blue, and phosphorylated residues are in red.

### Phosphorylation of axonemal dynein heavy chains in cytoplasm

We next used *Chlamydomonas* cytoplasmic extracts to ask whether any modifications identified in axonemal dynein HCs obtained from cilia are also present in the cytoplasmic pool of axonemal dyneins. We found multiple phosphorylated dynein peptides derived from the outer arm α HC and from six of the monomeric inner arm HCs (Table S3). These observations suggest that dyneins are subject to at least some phosphorylation events in the cytoplasm prior to their trafficking into the ciliary compartment. A total of ten phosphorylated sites (on the outer arm α HC and DHC2, DHC7, DHC8 and DHC9) were found in both cytoplasm and cilia, so potentially these additions occurred in the cytoplasm and the modified proteins were subsequently incorporated into the axoneme. In contrast, we identified five cytoplasmic phosphopeptides from the minor dyneins DHC4 and DHC11 that were not observed in our cilia-derived samples. Whether this reflects differential modification in the two cellular compartments or is a consequence of poor sequence coverage and depth due to the relatively low abundance of these HCs in the axoneme is uncertain.

### Stoichiometry of dynein heavy chain phosphorylation

To estimate the relative abundance of modification of each identified phosphorylation site, we examined the measured precursor intensities for the phosphopeptides and corresponding unmodified sequences identified by mass spectrometry (Fig. 7 and Table S4). The calculated stoichiometries exhibit a very broad range from <5% phosphorylated to numerous instances where the unmodified peptide was not found; in such cases, the stoichiometry was set as 100% modified. Use of *k*-means clustering of the phosphorylation stoichiometry values placed the modified sites into five groupings including those which are completely or almost completely modified, those modified at a level below ∼20%, and several clusters with intermediate levels of modification (∼60-80%, ∼40-60% and ∼20-40%). This suggests these various modifications might play distinct roles in dynein assembly (where complete modification might be expected) *versus* regulatory control.

**Figure 7.**
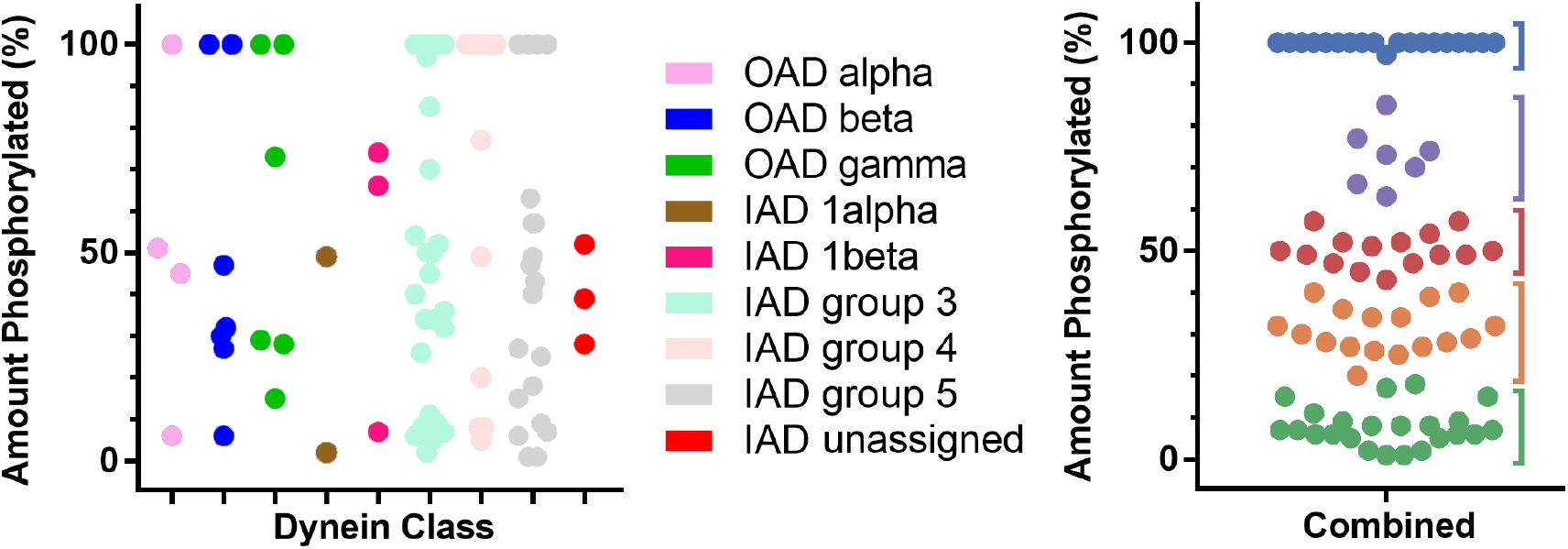
Phosphorylation Stoichiometry at Dynein Heavy Chain Sites. The *left* panel shows the percentage phosphorylation at every site identified in dynein HCs of each class. These data derive from the peptide intensities for the phosphorylated and unmodified forms measured during mass spectrometry analysis; the underlying intensity data are presented in Table S4. The *right* panel shows the data for all HCs combined as a single plot. These sites fall into apparent groupings based on *k*-means clustering of the modification stoichiometry values with *k* = 5. The identification of groups with such divergent modification stoichiometries suggests phosphorylation may play numerous and diverse roles in dynein assembly and regulation.

### Differential phosphorylation of *Chlamydomonas* dynein components in *cis* and *trans* cilia

The *Chlamydomonas* α HC has been implicated in the inherent beat frequency imbalance between the two cilia (Sakakibara et al., 1991). We observed in wildtype *Chlamydomonas* outer arm samples that the α HC contains two sites (S_3630_ and S_3657_) that are modified at a level of ∼50% which might reflect differences between these motors in the *cis* and *trans* cilia. To assess this, we prepared axonemal dynein HCs from the *Chlamydomonas uni1-1* (CC-2506) mutant which only assembles the *trans* cilium (Huang et al., 1982); unfortunately, no mutants assembling just the *cis* cilium exist. Following mass spectrometry of samples from the *uni1-1* mutant, we found sub-stoichiometric levels of phosphorylation at three sites (S_3657_, S_3667_ and S_3670_) but no modifications at S_3630_ and S_3674_ which were readily found in wildtype samples. Potentially then, one or both of these last two phosphorylated sites may be *cis* cilium-specific. We also examined HCs from *uni1-1* cytoplasm and did not find any phosphorylated α HC residues suggesting that these modifications occur in cilia.

The studies of (Takada and Kamiya, 1997) indicated that the outer arm docking complex is also involved in beat frequency imbalance again implying that it is somehow different in the two cilia. Examination of the phosphorylation stoichiometry for wildtype docking complex components based on the mass spectral peak intensities (Table S5) revealed that DC1 has several sites which appeared completely modified while others were at or below ∼25% phosphorylated. In contrast, DC2 had five sites that were ∼40-65% phosphorylated which might be consistent with distinct versions of this protein in the two cilia. Examination of these proteins from *uni1-1 trans* cilia (Table S5) found only two phosphorylated sites (S_347_ and T_652_) on DC1 neither of which had been observed previously. Why no modifications at the DC1 sites found to be fully phosphorylated in wildtype were not in observed in *uni1-1* is uncertain but implies there may be additional alterations in PTMs when only one cilium is assembled. For DC2, two sites (S_12_ and S_15_) that in wildtype have modification levels of ∼57% and 40%, respectively, were found to be completely phosphorylated in *uni1-1*; the unmodified form of this peptide was not found. The modification stoichiometry at a third DC2 site (S_266_) also doubled from ∼17% in wildtype to ∼36% in *uni1-1*. These observations are consistent with phosphorylation at these three DC2 sites being specific to the *trans* cilium (Fig. 8). Potentially, modification at other site(s) observed only in wildtype DC2 (such as T_20_, S_267_, T_512_) may reflect *cis* cilia-specific events.

**Figure 8.**
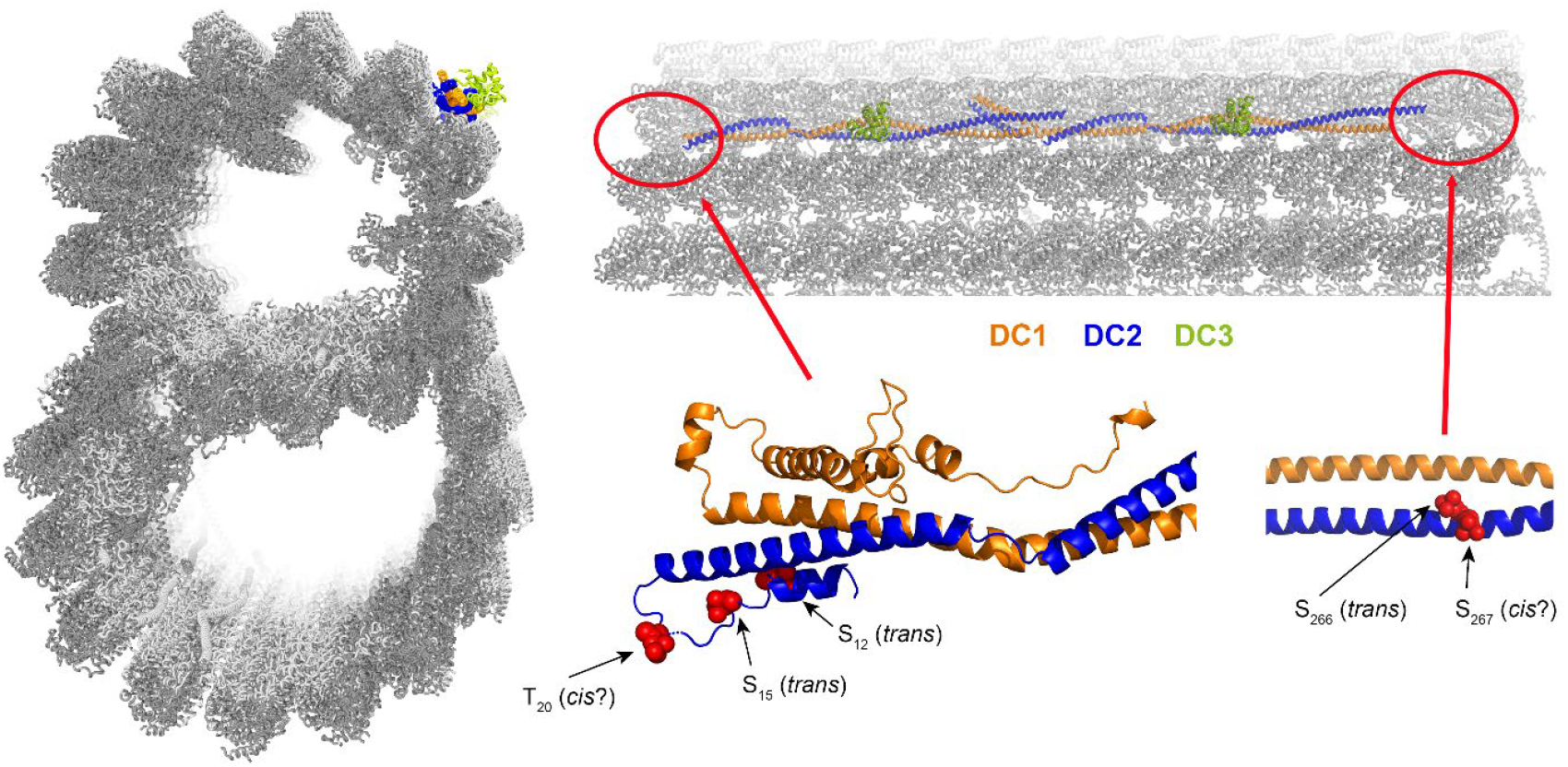
Cilium-specific Phosphorylation Sites on the Outer Arm Docking Complex. Two views of the cryo-EM reconstruction of the *Chlamydomonas* outer doublet microtubule highlighting the three-component outer arm docking complex (PDB 6U42); the color code is DC1 (orange), DC2 (blue) and DC3 (split-pea green). In the longitudinal view two docking complex repeats are shown. Large segments of DC1 and DC2 are missing from the structure, including the regions containing phosphorylated residues, presumably due to their inherent flexibility. At *lower right* are shown two segments of an AlphaFold 3 model of the docking complex illustrating the locations of five phosphorylated DC2 residues; their approximate locations on the *in situ* structure are indicated by red circles. Two phosphorylation sites identified with a stoichiometry near 50% in wildtype were found to be 100% modified in *uni1-1 trans* cilia. Modification at a third site (S_266_) essentially doubled in *uni1-1* although it remained substoichiometric. Also indicated are two other phosphorylation sites that were readily identified in wildtype samples but were missing from *uni1-1* suggesting they may be *cis*-cilium specific.

## Discussion

### A post-translational code for axonemal dynein heavy chains

Numerous systems are subject to PTMs that variably modify their properties and biological functions. Classic examples of these PTM codes are the methylation and acetylation of histones (Jenuwein and Allis, 2001), the numerous different modifications of tubulins that include acetylation, glycylation, glutamylation, methylation, (de)tyrosination, phosphorylation and succinylation (Janke and Magiera, 2020), and the alterations on actin isoforms such as acetylation, methylation, phosphorylation and sumoylation (Vedula and Kashina, 2018). Similarly, it is now becoming clear that dynein motors are also subject to various PTMs that might impact holoenzyme formation, axonemal incorporation and/or mechanochemical properties. Many of these motor enzyme proteins have processed N-termini whereby the terminal Met residue is either acetylated directly or removed by methionine aminopeptidase and the newly exposed N-terminal residue then acetylated (Sakato-Antoku et al., 2023). In addition, dyneins are methylated at numerous sites (Sakato-Antoku et al., 2024) and as discussed here are also subject to phosphorylation apparently by multiple kinases and dephosphorylation by one or more phosphatases.

All dynein HC phosphorylation modifications identified occur on Ser/Thr residues and we were not able to confirm any pTyr-containing phosphopeptides which is consistent with *in vivo* labeling and phosphorylated amino acid analysis in *Chlamydomonas* (King and Witman, 1994). That very distinct kinases controlled by different signaling pathways appear involved in dynein modifications provides the potential for signal integration at the level of individual arms. For example, analysis of the modified short linear motifs in various outer arm HCs revealed that PKC and CK II likely modify the β HC class, while PKA and unknown kinase(s) target the γ HC class. Furthermore, many of the modified regions are conserved in very distantly related eukaryotic supergroups. Consequently, although complex and difficult to immediately dissect, it is becoming clear that there may be a “dynein PTM code” that modifies individual motors. Examples of this include the recent observation that outer arm dynein stability is deficient in the absence of the DNAAF3/PF22 methyltransferase and that doublet microtubule-associated outer arms are extensively methylated while those trafficked into the cilium but remaining in the ciliary matrix are not (Sakato-Antoku et al., 2025). Dissecting this highly complex collection of dynein PTMs represents a major future experimental challenge.

### Binary classification of outer arm dyneins based on the α heavy chain

Most motile ciliated eukaryotes have both outer and inner rows of dynein arms. Exceptions include spermatozoids of bryophytes and vascular plants (such as ferns, cycads and *Gingko*) which lack outer arms, and diatoms (*e*.*g. Thalassiosira*) which do not encode inner arm dyneins (Wickstead, 2018). However, there is a further dichotomy based on the presence or absence of an ortholog of the outer arm α HC (Wickstead, 2018). For instance, this motor is missing from all Metazoa as well as the Metamonada (*e*.*g. Giardia*) and Discoba (*e*.*g. Trypanosoma*). In contrast, an α HC ortholog is present in motile members of the Chlorophyta, Cryptophyta and Haptophyta algae all of which have two cilia, as well as Stramenopila (*e*.*g*. diatoms) and Alveolata (*e*.*g. Tetrahymena, Paramecium* and dinoflagellates). All these latter organisms are single-celled and free-swimming in the environment, and it has been suggested previously that this HC type might be required for some types of tactic or environmental responses although it is dispensable at least for phototaxis in *Chlamydomonas* (Sakakibara et al., 1991).

This α HC class has several unusual structural features compared to its paralog the β HC. One defining characteristic is a very distinct N-terminal domain consisting of kelch and immunoglobulin repeats which form a β-propeller that associates with the remainder of the outer arm to form a HC trimer; the other two HCs interact *via* a dimerization interface that is generally conserved in all dimeric dynein HC motors. The α HC is also important for the retention of Lis1 in the cilium in response to treatments that impact normal ciliary beating (Rompolas et al., 2012) (discussed further below). Here we uncover a third unusual feature of this motor class – the phosphorylated extended loop structure that arches over the buttress coiled coil derived from AAA5.

### A role for phosphorylation in cis-trans frequency imbalance

In *Chlamydomonas*, the two cilia are not equivalent. The *trans* cilium, which derives from the mother basal body, has a higher inherent beat frequency than the *cis* cilium which emanates from the daughter basal body. This imbalance is lost in the *oda11* mutant which lacks the outer arm α HC suggesting that this motor is responsible for the differential activity and predictably is somehow different in the two cilia (Sakakibara et al., 1991). Furthermore, when wildtype dyneins are rebound to detergent-extracted cell models derived from the *oda1* mutant (lacks both outer arms and the outer arm docking complex) beat frequency imbalance is not restored rather the two cilia both beat at an intermediate frequency, but if rebound to *oda6* (lacks the outer arm but retains the docking complex) cell models the imbalance returns (Sakakibara and Kamiya, 1989; Takada and Kamiya, 1997). These observations indicate that the frequency imbalance is inherent to the axoneme and does not involve membrane-associated or ciliary matrix components. Furthermore, they imply that there is also a key essential role for the docking complex in controlling the inherent beat frequency of cilia. Together these data predict that outer arms in the two cilia are somehow different, perhaps with different PTMs, that might give rise to the imbalance which ultimately affects the ability of the cell to exhibit tactic responses.

To test whether phosphorylation might play a role in this process, we examined dyneins from the *uni1-1* mutant which only assembles a *trans* cilium. Although the α HC appears important for beat frequency imbalance, no clear *trans* cilia phosphorylated sites were observed in *uni1-1*. However, two sites readily found in wildtype were missing from *uni1-1* cilia so potentially these might represent *cis* cilia-specific sites; we are currently unable to test this concept further as no appropriate mutant exists. In contrast, we did find three sites on docking complex protein DC2 modified at levels consistent with their being *trans* cilia-specific. Analysis of the axonemal 96 nm repeat reconstruction (8GLV; (Walton et al., 2023)) reveals there is no direct contact between the outer arm α HC and the identified docking complex protein regions suggesting that any docking complex-derived signal is likely propagated through other outer arm components. However, most of the phosphorylation sites on DC2 occur in apparently disordered segments not included in the cryo-EM reconstructions and so their precise structural relationship to the rest of the outer arm remains uncertain. Additional evidence of a role for the docking complex in subtle control of ciliary motility comes from studies of the effects of alcohol on the phosphorylation of DC1, and the demonstration that three DC1 phosphorylated sites (S_33_, T_351_ and S_628_) are involved in the slowing of *Chlamydomonas* swimming velocity in response to alcohol treatments (Yang et al., 2019).

### Phosphorylation, Lis1 loading and increased force output in response to load

The dynein processivity factor Lis1 allows cytoplasmic dynein to transport large cargoes (*i*.*e*. nuclei) under high load conditions by slowing the rate of force-induced microtubule detachment (Kusakci et al., 2024). Lis1 also occurs in cilia (Pedersen et al., 2007) and the amounts present increase considerably in response to conditions that disrupt normal ciliary motility (Rompolas et al., 2012). This response is completely abrogated in the strain *oda11* (Rompolas et al., 2012) which lacks the outer arm α HC (Sakakibara et al., 1991); indeed, Lis1 was not detected in *oda11* cilia. In *Chlamydomonas*, treatments that reduce or inhibit beat frequency including viscous loading, alterations in the ciliary redox state and mutation of key systems required for motility such as the radial spokes and central pair apparatus all resulted in increased ciliary retention of Lis1. Furthermore, cells subject to high viscous load to enhance ciliary Lis1 levels and then returned to normal conditions exhibit an increased *trans* cilia beat frequency. This is consistent with higher force output by Lis1-bound outer arms in that cilium. In contrast, *cis* cilia on the same cells show little change in beat frequency (Rompolas et al., 2012). In cytoplasmic dynein, Lis1 associates with the AAA ring near AAA3 (DeSantis et al., 2017). Modeling of the *Chlamydomonas* α HC and Lis1 using AlphaFold 3 suggests that Lis1 docks very close to the flexible and extended phosphorylated region that loops over the AAA5-derived buttress. Consequently, it is feasible that differential alterations of this segment in the *cis* and *trans* cilia impact on whether or not Lis1 is retained during inhibition of ciliary beating.

### Extensive phosphorylation of inner dynein arms

Approximately 75% of the identified dynein HC phosphorylated sites occur in the monomeric inner arm HC groups. Furthermore, about two thirds of these sites map to within a few hundred residues of the HC N-termini which represents a very highly conserved post-translational feature of this dynein class that is found in some metazoans as well as trypanosomes, green alga, and alveolates. Modeling of groups IAD-3 and IAD-4 with and without the actin and p28/DNALI1 dimer that form part of these holoenzymes predict that all N-terminal sites are in unstructured loop regions that potentially interact with other axonemal components. Indeed, although the extreme N-terminus is missing, examination of the cryo-EM reconstruction of the *Chlamydomonas* 96-nm repeat (PDB 8GLV) reveals that the general region subject to phosphorylation is close to components of the calmodulin and spoke-associated complex (CSC) (Dymek et al., 2011; Walton et al., 2023). For IAD-5, modeling suggests the phosphorylated sites form two clusters between the HC and actin and centrin monomers. Thus, phosphorylation may be important for assembly of these dynein motors perhaps prior to axonemal incorporation.

In conclusion, we describe here the phosphorylation of axonemal dynein HCs from a broad array of motile ciliated eukaryotes. These extensive data suggest that phosphorylation of Ser/Thr residues apparently by numerous distinct kinases may play multiple roles in dynein assembly and/or have regulatory impacts on dynein-driven ciliary motility. These general processes and modified regions have been broadly retained across the eukaryotes providing a conservation-based guide to dissecting the functional role of phosphorylation on different dynein motor types.

## Materials and Methods

### Dynein heavy chain sources and preparation

Preparation of dynein HC-containing samples from twelve eukaryotic species and subsequent SDS-polyacrylamide electrophoresis was described in detail previously (Sakato-Antoku et al., 2025). The dynein sources examined were purified outer arm dynein from *Tetrahymena thermophila*, comb plates from *Mnemiopsis leidyi* (ctenophore), sperm axonemes from *Ciona intestinalis* (sea squirt), *Crassostrea gigas* (Pacific oyster), *Hemicentrotus pulcherrimus* (sea urchin), *Oncorhynchus mykiss* (rainbow trout), and *Takifugu rubripes* (puffer), cilia from *Chlamydomonas reinhardtii* (green alga), flagella from *Trypanosoma brucei* (trypanosome), seminal vesicles from *Drosophila melanogaster* (fruit fly) and *Drosophila willistoni* (fruit fly), and spermatozoids from *Ceratopteris richardii* (water fern). SDS polyacrylamide gels were stained with Coomassie blue, and the dynein HC-containing regions excised and processed for mass spectrometric analysis.

We also examined phosphorylation in whole cilia samples from *Chlamydomonas reinhardtii* wildtype (CC-125) cells and both the HC and 50-150 kDa regions of gels loaded with cilia from the *uni1-1* (CC-2506) mutant which only assembles *trans* cilia. These strains were grown, deciliated and the cilia subsequently isolated and processed using our standard methods described in detail in (Sakato-Antoku and King, 2023).

### Mass spectrometry

Dynein HC and cilia samples were digested with trypsin and/or endoproteinase Asp-N, peptides isolated, and mass spectrometry performed using a Orbitrap Eclipse Tribrid mass spectrometer using a Dionex Ultimate 3000 RSLCnano UPLC system with a 75 μm x 25 cm nanoEase m/z peptide BEH C18 column as described in detail in (Sakato-Antoku et al., 2024; Sakato-Antoku et al., 2025). Raw data were searched using the Andromeda search engine embedded within MaxQuant. All the organismal proteomes analyzed have been made available in fasta format at the ProteomeXchange Consortium *via* the PRIDE partner repository associated with the dataset identifiers PXD055793 and PXD058873. Mass spectral searches included the following variable modifications: Met oxidation, protein N-terminal acetylation, methyl Lys/Arg, dimethyl Lys/Arg, trimethyl Lys/Arg and Ser/Thr/Tyr phosphorylation; carbamidomethylation of Cys was included as a fixed modification. The searches of raw data associated with the *uni1-1* mutant did not include methylation of Arg and Lys residues. Data were visualized using Scaffold v.5.0.1 (Proteome Software).

### Protein structure modeling and display

Models structures for regions of various dynein HCs and their binding partners including added nucleotides, metal ions and phosphorylated residues as appropriate were generated using AlphaFold 2 and AlphaFold 3 (Abramson et al., 2024; Jumper et al., 2021). When modeling the entire AAA ring of HCs, the N-terminal regions were truncated due to size limitations imposed by the AlphaFold 3 server. Structures were displayed using the PyMOL molecular graphics system (Schrödinger, LLC). Structures were superimposed using the commands “align” and “super” within PyMOL.

## Bioinformatics

Phosphorylated sequence motifs were aligned manually, and consensus sequences displayed using WebLogo. Sequences phosphorylated by unknown kinases were analyzed using the Kinase Predictor v.0.8 tool which may be downloaded from esbl.nhlbi.nih.gov/Databases/Kinase_Logos/KinasePredictor.html.

All figures were prepared using Adobe Illustrator and/or Photoshop.

## Abbreviations

HC: heavy chain
IC: intermediate chain
LC: light chain
PTM: post-translational modification.

## Nomenclature

Dynein heavy chain nomenclature is confusing as often the same name has been assigned to non-orthologous sequences in different species. Consequently, here we utilize the conventions from *Chlamydomonas*. See (Braschi et al., 2022; Hom et al., 2011) for other naming schemes and heavy chain equivalence.

## Funding

This study was supported by grant R35-GM140631 from the National Institutes of Health (to SMK).

## Author Contributions

Conceptualization, SMK; Methodology, JLB, SMK; Investigation and Sample Preparation, MS-A, RSP-K, KI, JLB; Visualization, JLB, SMK; Supervision, SMK; Writing – Original Draft, SMK, JLB; Writing – Review and Editing, MS-A, RSP-K, KI, JLB, SMK.

## Competing Interests

The authors declare no competing interests.

## Acknowledgements

We thank Dr Adam Schuyler (UConn Health) for performing the *k*-means clustering analysis and Drs Ramaswamy Chidambaram (UConn Health), Nikisha Patel (Trinity College), Mayu Inaba (UConn Health), Qinhui Rao (Yale), Jun Yang (Yale), Mark Terasaki (UConn Health), Michelle Shinogawa (UCLA) and Kent Hill (UCLA) for providing samples for our previously published and released dynein heavy chain mass spectral datasets. We acknowledge the NIH S10 high-end instrumentation award 1S10-OD028445-01A1, which supported this work by providing funds to acquire the Orbitrap Eclipse Tribrid mass spectrometer housed in the University of Connecticut Proteomics and Metabolomics Facility.

## Data Availability

The mass spectrometry data for *Chlamydomonas* wildtype samples are available at Dryad with the identifiers fn2z34txn and mw6m90635 and at PRIDE with the dataset identifier PXD045935. Raw mass spectral data and the searched proteomes (in fasta format) for all other organisms have been deposited to the ProteomeXchange Consortium *via* the PRIDE partner repository with the dataset identifiers PXD055793 and PXD058873. The *Chlamydomonas uni1-1* datasets have been deposited *via* PRIDE with the identifier PXD061468.

## Data Access for Reviewers

The datasets fn2z34txn and mw6m90635 are publicly available at Dryad (doi: fn2z34txn and doi: mw6m90635)

The PXD055793 and PXD058873 datasets are publicly available at PRIDE (https://www.ebi.ac.uk/pride/)

The PXD061468 dataset may be accessed with the following log-on credentials:

**Password:** 9LBcMI1HFAIX

## Notes

### Competing Interest Statement

The authors have declared no competing interest.

