## Supplemental Data for "Isoform-specific Patterns of Phosphorylation on Axonemal Dynein Heavy Chains"

**and Stephen M. King**

| | |
|---|---|
| Figure S1 | Putative Site of Lis1 Interaction Near the Phospho-Loop of the Outer Arm $\alpha$ Heavy Chain |
| Table S1 | Phosphorylated Sites on Dynein Heavy Chains Identified by Mass Spectrometry |
| Table S2 | Phosphorylated Dynein Heavy Chain Site Matches to Kinase Consensus Sequences |
| Table S3 | Phosphorylated Sites on *Chlamydomonas* Dynein Heavy Chains Identified in Cytoplasm |
| Table S4 | Modification Stoichiometry at Dynein Heavy Chain Phosphorylation Sites Based on Mass Spectral Peak Intensities |
| Table S5 | Modification Stoichiometry at Outer Arm Docking Complex Phosphorylation Sites in Wildtype (CC-125) and *uni1-1* Mutant Cilia Based on Mass Spectral Peak Intensities |

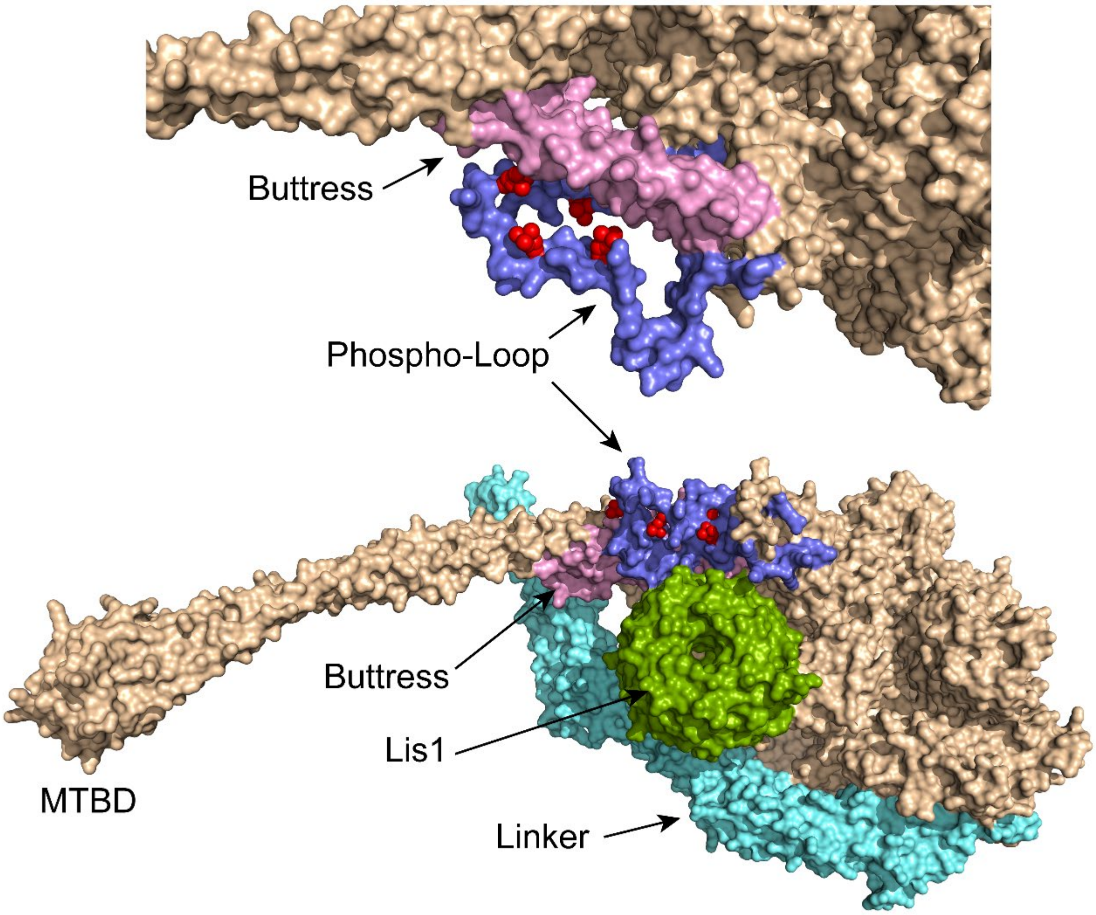


**Figure S1 Putative Site of Lis1 Interaction Near the Phospho-Loop of the Outer Arm α Heavy Chain**

The *upper* panel is an AlphaFold 3 model of the α HC region including the buttress and phospho-loop. Three of the phosphoryl groups are oriented towards the outward coil of the buttress. The *lower* panel shows the predicted location of Lis1 adjacent to the buttress and phospho-loop. The color code is light pink (buttress), cyan (lever arm with N-terminal β-propeller region truncated), slate blue (phospho-loop), red (phosphorylated residues), split pea green (Lis1). The α HC and Lis1 sequences are from Phytozome accessions Cre03.g145127 and Cre12.g552900, respectively.

**Table S1** **Phosphorylated Sites on Dynein Heavy Chains Identified by Mass Spectrometry***

| Organism | Source | Dynein Heavy Chain Designation | Residues x/n (%) | Sequence Coverage MS/MS Spectra Total | Unique | Peptides Unique | Phosphorylated | Phosphorylated Residues |
|---|---|---|---|---|---|---|---|---|
| ***Outer Arm $\alpha$ HC*** | | | | | | | | |
| *Chlamydomonas* | Cilia | DHC13 | 3441/4503 (76%) | 1131 | 553 | 311 | 7 | $S_{3630}$, $S_{3657}$, $S_{3674}$ (Pan et al. 2011) $S_{3649}$, $S_{3670}$ (Wang et al. 2014) $S_{3667}$ |
| *Tetrahymena* | | TTHERM_00486600 DYH5 ($\gamma$ HC) | 3075/4168 (74%) | 1112 | 479 | 314 | 3 | $S_{3381}$ |
| ***Outer Arm $\beta$ HC*** | | | | | | | | |
| *Chlamydomonas* | Cilia | DHC14 | 3771/4568 (83%) | 1251 | 633 | 366 | 1 | $T_{60}$ |
| *Ciona* | Sperm Flagella | Ci_11920 DNAH9 C-terminal region missing | 2061/3522 (59%) | 645 | 266 | 191 | 0 | --- |
| *Crassostrea* | Sperm Flagella | XP_011428027 DNAH9 | 2768/4465 (62%) | 887 | 412 | 302 | 3 | $S_3$ |
| *Drosophila* | Seminal Vesicles #1 | Kl-5 | 1986/4559 (44%) | 339 | 211 | 185 | 0 | --- |
| | Seminal Vesicles #2 | Kl-5 | 2244/4559 (49%) | 503 | 317 | 228 | 0 | --- |
| *Hemicentrotus* | Sperm Flagella | HPU_11785 $\beta$ HC | 2465/4076 (60%) | 805 | 369 | 251 | 0 | --- |
| *Mnemiopsis* | Comb Plates | ML002216a partial | 1063/3176 (34%) | 147 | 129 | 111 | 0 | --- |
| *Oncorhynchus* | Sperm Axonemes | DNAH17 | 1993/4460 (45%) | 605 | 286 | 227 | 0 | --- |
| *Rattus* | Tracheal Cilia | DNAH9 | 2850/4410 (65%) | 972 | 521 | 295 | 1 | $S_{13}$ |
| | | DNAH11 | 2748/4487 | 702 | 410 | 269 | 3 | $S_{13}$ |

| | | | | | | | | |
|---|---|---|---|---|---|---|---|---|
| | | | (61%) | | | | | |
| | | DNAH17 | 240/4459 (5%) | 84 | 1 | 1 | 0 | --- |
| *Tetrahymena* | Purified Dynein | TTHERM_00499300 DYH4 | 3384/4595 (74%) | 1284 | 562 | 380 | 0 | --- |
| *Trypanosoma* | Flagella | Tb927.11.3250 | 3747/4658 (80%) | 1461 | 791 | 412 | 4 | $S_{1050}$, $S_{1056}$, $S_{4089}$, $S_{4091}$ |
| ***Outer Arm γ HC*** | | | | | | | | |
| *Chlamydomonas* | Cilia | DHC15 | 3590/4501 (80%) | 1250 | 621 | 358 | 0 | --- |
| *Ciona* | Sperm Flagella | Ci_18292 DNAH8 | 2634/4239 (62%) | 991 | 390 | 266 | 4 | $S_{876}$, $S_{971}$ |
| *Crassostrea* | Sperm Flagella | XP_034301736 DNAH8 | 2933/4648 (63%) | 950 | 391 | 283 | 2 | $S_{29}$, $S_{48}$ |
| *Drosophila* | Seminal Vesicles #1 | Kl-3 | 1560/4593 (34%) | 248 | 176 | 160 | 0 | --- |
| | Seminal Vesicles #2 | Kl-3 | 2019/4593 (46%) | 434 | 278 | 210 | 0 | --- |
| *Hemicentrotus* | Sperm Flagella | HPU_07454[@] α HC DNAH8 | 1314/2408 (55%) | 385 | 163 | 117 | 5 | $T_{1071}$, $S_{1092}$ (in *S. purpuratus*) |
| | | HPU_14295[@] α HC DNAH5 | 166/2265 (7%) | 54 | 8 | 6 | 0 | --- |
| *Mnemiopsis* | Comb Plates | ML07114a | 1420/4505 (32%) | 192 | 156 | 145 | 0 | --- |
| *Oncorhynchus* | Sperm Axonemes | DNAH8 | 2049/4362 (47%) | 523 | 264 | 215 | 0 | --- |
| *Rattus* | Tracheal Cilia | DNAH5 | 2798/4621 (61%) | 975 | 512 | 293 | 0 | --- |
| *Tetrahymena* | Purified Dynein | TTHERM_01276420 DYH3 | 3701/4620 (80%) | 1371 | 620 | 412 | 0 | --- |
| *Trypanosoma* | Flagella | Tb927.3.930 | 3595/4639 (77%) | 1498 | 811 | 410 | 0 | --- |
| ***Inner Arm I1/f 1α HC*** | | | | | | | | |

| | | | | | | | | |
|---|---|---|---|---|---|---|---|---|
| *Ceratopteris* | Spermato-zoids #1 | Ceric_25G055400 | 177/4556 (4%) | 16 | 15 | 15 | 0 | --- |
| | Spermato-zoids #2 | Ceric_25G055400 | 483/4556 (11%) | 56 | 48 | 47 | 0 | --- |
| *Chlamydomonas* | Cilia | DHC1 | 3232/4626 (70%) | 562 | 398 | 279 | 1 | $S_{1050}$ |
| *Crassostrea* | Sperm Flagella | XP_034298943 DNAH10 | 2596/4668 (56%) | 522 | 328 | 260 | 1 | $S_{111}$ |
| *Drosophila* | Seminal Vesicles #1 | DHC98D | 942/5080 (19%) | 107 | 91 | 86 | 0 | --- |
| | Seminal Vesicles #2 | DHC98D | 1432/5080 (28%) | 184 | 157 | 134 | 0 | --- |
| *Hemicentrotus* | Sperm Flagella | HPU_17685 DNAH10 partial | 1236/2469 (50%) | 255 | 158 | 128 | 0 | --- |
| *Mnemiopsis* | Comb Plates | ML03391a | 504/2564 (20%) | 57 | 54 | 52 | 0 | --- |
| *Oncorhynchus* | Sperm Axonemes | DNAH10 | 1607/4629 (35%) | 249 | 179 | 160 | 0 | --- |
| *Rattus* | Tracheal Cilia | DNAH10 | 2550/4592 (56%) | 594 | 393 | 262 | 0 | --- |
| *Takifugu* | Sperm Axonemes #1 | Tr_742129 DNAH10 | 1349/4228 (32%) | 218 | 107 | 91 | 0 | --- |
| | Sperm Axonemes #2 | Tr_742129 DNAH10 | 2271/4228 (54%) | 576 | 287 | 167 | 0 | --- |
| *Tetrahymena* | Purified Dynein | TTHERM_00688470 DYH6 | 3037/4383 (69%) | 704 | 420 | 313 | 0 | --- |
| *Trypanosoma* | Flagella | Tb927.4.870 | 3384/4599 (74%) | 756 | 561 | 322 | 0 | --- |
| ***Inner Arm I1/f 1β HC*** | | | | | | | | |
| *Ceratopteris* | Spermato-zoids #1 | Ceric_22G060000 partial | 125/2814 (4%) | 12 | 10 | 10 | 0 | --- |

| | | | | | | | | |
|---|---|---|---|---|---|---|---|---|
| | Spermato-zoids #2 | Ceric_22G060000 partial | 284/2814 (10%) | 34 | 28 | 28 | 0 | --- |
| *Chlamydomonas* | Cilia | DHC10 | 3315/4513 (73%) | 615 | 428 | 284 | 0 | --- |
| *Crassostrea* | Sperm Flagella | XP_034330723 DNAH2 | 2422/4554 (60%) | 474 | 308 | 239 | 0 | --- |
| *Drosophila* | Seminal Vesicles #1 | Kl-2 | 815/4459 (18%) | 107 | 79 | 77 | 0 | --- |
| | Seminal Vesicles #2 | Kl-2 | 1113/4459 (25%) | 146 | 125 | 110 | 0 | --- |
| *Hemicentrotus* | Sperm Flagella | HPU_13585 DNAH2 | 1818/3780 (48%) | 362 | 220 | 167 | 2 | $S_{42}$ |
| *Mnemiopsis* | Comb Plates | ML14857a partial | 430/2191 (20%) | 48 | 41 | 41 | 0 | --- |
| *Oncorhynchus* | Sperm Axonemes | DNAH2 | 1221/3930 (31%) | 172 | 125 | 109 | 0 | --- |
| *Rattus* | Tracheal Cilia | DNAH2 | 2388/4508 (53%) | 500 | 359 | 228 | 0 | --- |
| *Takifugu* | Sperm Axonemes #1 | Tr_732194 DNAH2 | 1115/4685 (24%) | 159 | 125 | 114 | 0 | --- |
| | Sperm Axonemes #2 | Tr_732194 DNAH2 | 2269/4685 (48%) | 575 | 410 | 234 | 1 | $S_{125}$ |
| *Tetrahymena* | Purified Dynein | TTHERM_00912290 DYH7 | 3066/4805 (64%) | 698 | 434 | 309 | 1 | $T_{3229}$ |
| *Trypanosoma* | Flagella | Tb927.8.3250 | 3289/4674 (70%) | 707 | 527 | 317 | 0 | --- |
| ***Inner Arm Group 3 HCs*** | | | | | | | | |
| *Ceratopteris* | Spermato-zoids #1 | Ceric_20G075500 | 111/4153 (3%) | 12 | 9 | 9 | 0 | --- |
| | | Ceric_03G083100) | 120/4178 (3%) | 12 | 9 | 9 | 0 | --- |

| | | | | | | | | |
|---|---|---|---|---|---|---|---|---|
| | Spermato-zoids #2 | Ceric_20G075500 | 456/4153 (11%) | 51 | 47 | 46 | 0 | --- |
| | | Ceric_03G083100 | 459/4178 (11%) | 53 | 44 | 44 | 0 | --- |
| *Chlamydomonas* | Cilia | DHC4 | 516/4906 (11%) | 49 | 28 | 28 | 3 | $T_{941}$, $S_{945}$, $S_{4532}$ |
| | | DHC5 | 2484/4185 (59%) | 414 | 271 | 179 | 0 | --- |
| | | DHC6 | 2651/4006 (66%) | 369 | 285 | 200 | 2 | $T_{128}$, $T_{205}$<br>(Wang et al. 2014) $S_{45}$, $S_{46}$<br>(Wagner et al 2006) $S_{3435}$, $S_{3450}$, $S_{3452}$ |
| | | DHC8 | 2714/4209 (64%) | 434 | 307 | 217 | 3 | $S_{88}$<br>(Wang et al. 2014) $T_{33}$, $S_{35}$<br>(Pan et al. 2011) $T_{76}$, $S_{1960}$, $T_{1961}$ |
| | | DHC9 | 2993/4152 (72%) | 589 | 405 | 252 | 4 | $S_{155}$, $S_{159}$ |
| | | DHC11 | 497/4757 (10%) | 44 | 29 | 28 | 0 | --- |
| *Crassostrea* | Sperm Flagella | XP_034301924 DNAH7 | 2122/4022 (53%) | 477 | n/a† | n/a† | 2 | $T_{142}$ |
| | | XP_034306561 DNAH12 | 2014/4030 (50%) | 352 | n/a† | n/a† | 0 | --- |
| | | XP_011412296 DNAH3 | 1850/4073 (45%) | 343 | 211 | 168 | 4 | $T_{11}$, $S_{15}$, $S_{107}$ |
| *Drosophila* | Seminal Vesicles #1 | DNAH3 | 806/4385 (18%) | 94 | 73 | 70 | 0 | --- |
| | | DHC36C | 958/4024 (24%) | 120 | 98 | 91 | 0 | --- |
| | | DHC62B | 767/3964 (19%) | 87 | 69 | 66 | 0 | --- |
| | Seminal Vesicles #2 | DNAH3 | 1514/4385 (35%) | 213 | 169 | 141 | 0 | --- |
| | | DHC36C | 1683/4024 (42%) | 298 | 215 | 159 | 0 | --- |

| | | | | | | | | |
|---|---|---|---|---|---|---|---|---|
| | | DHC62B | 1332/3964 (34%) | 188 | 148 | 120 | 0 | --- |
| *Hemicentrotus* | Sperm Flagella | HPU_12543 DNAH7 | 2352/3926 (60%) | 564 | 302 | 206 | 1 | $S_{147}$ |
| | | HPU_18186[+] DNAH12 | 1911/3683 (52%) | 369 | 233 | 173 | 3 | $S_{57}$, $T_{1593}$, $T_{3405}$ |
| *Mnemiopsis* | Comb Plates | ML23952a | 636/4054 (16%) | 64 | 55 | 55 | 0 | --- |
| | | ML329912a | 615/3846 (16%) | 63 | 55 | 53 | 0 | --- |
| *Oncorhynchus* | Sperm Axonemes | DNAH3 | 1412/4063 (35%) | 236 | 154 | 130 | 4 | $S_{126}$, $T_{169}$ |
| | | DNAH7 | 1549/4013 (39%) | 290 | 172 | 140 | 3 | $S_{70}$, $S_{85}$, $S_{144}$ |
| | | DNAH12 | 1263/3964 (32%) | 219 | 145 | 125 | 0 | --- |
| *Rattus* | Tracheal Cilia | DNAH3 | 1910/4069 (47%) | 366 | 245 | 169 | 0 | --- |
| | | DNAH7 | 2677/4023 (67%) | 700 | 419 | 243 | 2 | $T_{152}$ |
| | | DNAH12 | 2086/3960 (53%) | 407 | 279 | 196 | 0 | --- |
| *Takifugu* | Sperm Axonemes #1 | Tr_188439 DNAH3 | 1991/4013 (50%) | 841 | 303 | 222 | 6 | $S_{78}$, $S_{120}$ |
| | | Tr_728981 DNAH12 | 1246/3581 (35%) | 203 | 140 | 120 | 0 | --- |
| | Sperm Axonemes #2 | Tr_188439 DNAH3 | 3023/4013 (75%) | 2031 | 752 | 343 | 15 | $S_{68}$, $S_{78}$ |
| | | Tr_728981 DNAH12 | 2076/3581 (58%) | 540 | 365 | 203 | 0 | --- |
| *Tetrahymena* | Purified Dynein | TTHERM_00252430 DYH11 | 997/4257 (23%) | 101 | 77 | 72 | 0 | --- |

| | | | | | | | | |
|---|---|---|---|---|---|---|---|---|
| | | TTHERM_00774810 DYH25 (not full-length) | 958/2425 (40%) | 104 | 87 | 73 | 1 | $T_{1092}$ |
| *Trypanosoma* | Flagella | Tb927.2.5270 | 3241/4246 (76%) | 700 | 542 | 309 | 1 | $S_{152}$ |
| ***Inner Arm Group 4 HCs*** | | | | | | | | |
| *Ceratopteris* | Spermato-zoids #1 | Ceric_25G028800 | 311/4052 (8%) | 31 | 27 | 27 | 0 | --- |
| | Spermato-zoids #2 | Ceric_25G028800 | 646/4052 (16%) | 74 | 69 | 65 | 0 | --- |
| *Chlamydomonas* | Cilia | DHC2 | 2236/3478 (64%) | 446 | 285 | 187 | 5 | $S_{52}$, $S_{192}$, $S_{2480}$, $T_{2482}$ |
| *Crassostrea* | Sperm Flagella | XP_034334098 DNAH1 | 2212/4211 (52%) | 433 | 278 | 212 | 2 | $S_{9}$, $S_{16}$ |
| *Mnemiopsis* | Comb Plates | ML053015a | 993/3999 (25%) | 110 | 98 | 94 | 0 | --- |
| | | ML34752a partial | 33/1077 (3%) | 3 | 3 | 3 | 0 | --- |
| *Oncorhynchus* | Sperm Axonemes | DNAH1 | 1672/4208 (40%) | 269 | 184 | 153 | 1 | $S_{100}$ |
| *Rattus* | Tracheal Cilia | DNAH1 | 2526/4250 (59%) | 516 | 338 | 231 | 2 | $S_{65}$ |
| *Tetrahymena* | Purified Dynein | TTHERM_001151438 DYH21 | 1815/3911 (46%) | 216 | 159 | 135 | 0 | --- |
| *Trypanosoma* | Flagella | Tb11.10.5350 | 3033/4142 (73%) | 623 | 480 | 293 | 2 | $S_{26}$ |
| | | Tb927.11.8160 | 2935/4152 (71%) | 620 | 458 | 269 | 3 | $S_{49}$ |
| ***Inner Arm Group 5 HCs*** | | | | | | | | |
| *Ceratopteris* | Spermato-zoids #1 | Ceric_29G018300 partial | 176/3375 (5%) | 20 | 15 | 15 | 0 | --- |
| | Spermato-zoids #2 | Ceric_29G018300 partial | 320/3375 (9%) | 40 | 34 | 33 | 0 | --- |
| *Chlamydomonas* | Cilia | DHC3 | 409/5593 | 38 | 24 | 24 | 4 | $S_{152}$, $S_{171}$, $S_{685}$, $S_{5326}$ |

| | | | | | | | | |
|---|---|---|---|---|---|---|---|---|
| | | | (7%) | | | | | (Wang et al. 2014) $S_{4265}$, $S_{4267}$ |
| | | DHC7 | 2914/4191 (70%) | 515 | 363 | 246 | 12 | $S_{9}$, $S_{90}$, $T_{104}$, $S_{111}$, $S_{660}$ (Wang et al 2014) $T_{87}$, $T_{93}$, $S_{120}$ |
| *Crassostrea* | Sperm Flagella | XP_034321313 DNAH6 | 2148/4220 (51%) | 394 | n/a | n/a | 3 | $S_{137}$, $S_{146}$ |
| *Drosophila* | Seminal Vesicles #1 | DHC16F | 967/4981 (24%) | 113 | 95 | 93 | 0 | --- |
| | Seminal Vesicles #2 | DHC16F | 1428/4081 (35%) | 205 | 167 | 132 | 0 | --- |
| *Hemicentrotus* | Sperm Flagella | HPU_06524 DNAH6 | 2101/4236 (50%) | 387 | 256 | 193 | 4 | $S_{19}$, $S_{1390}$, $T_{1400}$ |
| *Mnemiopsis* | Comb Plates | ML011724a partial | 139/900 (15%) | 18 | 15 | 14 | 0 | --- |
| *Oncorhynchus* | Sperm Axonemes | DNAH6 | 1094/3611 (30%) | 195 | 110 | 95 | 0 | --- |
| | | DNAH14 | 237/4450 (5%) | 30 | 20 | 20 | 0 | --- |
| *Rattus* | Tracheal Cilia | DNAH6 | 2311/4144 (56%) | 549 | 356 | 231 | 0 | --- |
| *Tetrahymena* | Purified Dynein | TTHERM_00193520 DYH24 | 1036/4126 (25%) | 108 | 85 | 74 | 0 | --- |
| | | TTHERM_00565600 DYH22 | 506/4564 (11%) | 58 | 36 | 35 | 0 | --- |
| *Trypanosoma* | Flagella | Tb927.11.11220 | 3096/4242 (73%) | 712 | 503 | 277 | 4 | $T_{155}$, $S_{158}$, $S_{222}$, $S_{3173}$ |
| | | Tb927.7.920 | 2922/4112 (71%) | 641 | 472 | 258 | 5 | $S_{7}$, $S_{18}$, $S_{75}$ |
| ***Unassigned Inner Arm HC*** | | | | | | | | |
| *Chlamydomonas* | Cilia | DHC12 | 247/6150 (4%) | 22 | 16 | 16 | 4 | $S_{758}$, $S_{1124}$, $S_{4848}$ (Wang et al 2014) $S_{168}$, $S_{297}$, $S_{299}$, $S_{301}$, $S_{334}$, $S_{2610}$, $S_{2612}$, $S_{3603}$, $S_{4951}$ |
| ***IFT Dynein HC*** | | | | | | | | |
| *Chlamydomonas* | Cilia | DHC16 | 3078/4333 (71%) | 615 | 426 | 291 | 0 | --- |

| | | | | | | | | |
|---|---|---|---|---|---|---|---|---|
| *Mnemiopsis* | Comb Plates | ML019112a partial | 22/2556 (1%) | 2 | 2 | 2 | 0 | --- |
| *Rattus* | Tracheal Cilia | DYNC2H1 | 2550/4306 (59%) | 434 | 320 | 241 | 0 | --- |
| *Trypanosoma* | Flagella | Tb927.4.560 | 25/4232 (1%) | 2 | 2 | 2 | 0 | --- |
| ***Cytoplasmic Dynein HC*** | | | | | | | | |
| *Drosophila* | Seminal Vesicles #1 | DHC64C | 850/4638 (18%) | 83 | 74 | 73 | 0 | --- |
| | Seminal Vesicles #2 | DHC64C | 1592/4638 (34%) | 182 | n/a | 138 | 0 | --- |
| *Rattus* | Tracheal Cilia | DYNC1H1 | 2441/4646 (53%) | 407 | 328 | 236 | 1 | $S_{634}$ |

* The raw mass spectral data from these samples were previously analyzed for the presence of methylated Arg and Lys residues. Consequently, the general sequence coverage is as presented in Sakato-Antoku *et al*. (2025).

@ The available *Hemicentrotus* sequences are incomplete. All phosphorylated basic residues identified in *Hemicentrotus* are conserved in the sea urchin *Strongylocentrotus purpuratus* and therefore the residue numbers from the equivalent heavy chain of that organism are shown.

+ The available *Hemicentrotus* sequence for DNAH12 is missing ~150 residues including the Walker A and B motifs of AAA1.

† Due to a very high degree of shared primary sequence for two closely related isoforms, we are unable to definitively assign spectra and peptides to individual proteins.

**Table S2** Phosphorylated Dynein Heavy Chain Site Matches to Kinase Consensus Sequences

| Organism | Sequence | Kinase Consensus Sequences* | | | | | | | Uncertain |
|---|---|---|---|---|---|---|---|---|---|
| | | cAMP- or cGMP-dependent Kinase | Protein Kinase C | Caesin Kinase II | $Ca^{2+}$/Calmodulin-dependent Protein Kinase II | Polo-like Kinase 1 | Mitogen-activated Protein Kinase | NimA-related Kinase (NEK) | |
| | $X_6$-[pS/pT]-$X_6$ | R[R/K]x[S/T] | [S/T]x[R/K] | [S/T]xx[D/E] | Rxx[S/T]x | [D/E/N]x[S/T] | [P/$\phi$]x[S/T]P | [L/F/M/W]xx[S/T][not P] | |
| ***Outer Arm $\alpha$ HC*** | | | | | | | | | |
| *Chlamydomonas* | GLKKTP**S**LRDQPM | | Yes | Yes | | | | | |
| | EKARRS**S**GVGDRR | Yes | | | | | | | |
| | VGDRRP**S**QEGLPG | Yes | | | | | | | |
| | LPGPEA**S**QASLAE | | | | | | | | Yes |
| | PEASQA**S**LAESQG | | | Yes | | | | | |
| | QASLAE**S**QGGRGS | | | | | | | Yes | |
| *Tetrahymena* | DSDEPM**S**PRSLKK | | Yes | | | | | | |
| ***Outer Arm $\beta$ HC*** | | | | | | | | | |
| *Chlamydomonas* | FMEEAH**T**KRLLIM | | Yes | | | | | | |
| *Crassostrea* | ----MA**S**ELELND |  |  | Yes |  |  |  |  |  |
| *Rattus* | QAVLAE**S**GDEEPG | | | Yes | | | | | |
| | GTVLRL**S**PSEQED | | | Yes | | | | | |
| *Trypanosoma* | VTKFHS**S**LTDEES | | | Yes | | | | | |
| | SLTDEE**S**VKEVAP | | Yes | Yes | | | | | |
| | QKVMED**S**GSEKAT | | | Yes | | | | | |
| | VMEDSG**S**EKATSL | | Yes | | | | | | |
| ***Outer Arm $\gamma$ HC*** | | | | | | | | | |
| *Ciona* | AYGRRM**S**DQLTNA | Yes | | | | | | | |
| | DGKKMG**S**QMTLQN | | | | | | | | Yes |
| *Crassostrea* | FYERRE**S**VIYANP | Yes | | | | | | | |
| | PGGRRR**S**TVSSVL | Yes | | | | | | | |
| *Hemicentrotus* | QWGQSR**T**RPNEPA | | | | | | | | Yes |
| | HRHGAH**S**APHVEV | | | | | | | | Yes |

| | | | | | | | | | |
|---|---|---|---|---|---|---|---|---|---|
| ***Inner Arm I1/f 1α HC*** | | | | | | | | | |
| *Chlamydomonas* | VRSLVE**S**TKAFVR | | Yes | | | | | | |
| *Crassostrea* | QPLTVS**S**PTEEEE | | | Yes | | | | | |
| ***Inner Arm I1/f 1β HC*** | | | | | | | | | |
| *Hemicentrotus* | NGGKSE**S**PPAIKL | | | | | | | | Yes |
| *Takifugu* | TEELTE**S**EKRQLN | | Yes | | | | | Yes | |
| *Tetrahymena* | PLKQEP**T**PVPVPE | | | | | Yes | | | |
| ***Inner Arm Group 3 HCs*** | | | | | | | | | |
| *Chlamydomonas* | VTFEPR**T**HRVSED | | Yes | | | | | | |
| | PRTHRV**S**EDGPAA | | | | | | | | Yes |
| | LADEDG**S**EGGGHD | | | | | | | | Yes |
| | LFESAL**S**SGNASL | | | | | | | | Yes |
| | FESALS**S**GNASLV | | | | | | | | Yes |
| | PPLPTA**T**PPVPRR | | | | | | | | Yes |
| | TRLRPR**T**GQLPVR | | | | Yes | | | | |
| | IEVLSA**S**EGNILE | | | | | | | Yes | |
| | TAINVI**S**SSKLLS | | Yes | | | | | | |
| | INVISS**S**KLLSND | | | | | | | | Yes |
| | GRLPGR**T**GSPGAG | | | | | | | | Yes |
| | LPGRTG**S**PGAGFT | | | | Yes | | | | |
| | EHLEFG**T**KRKKSA | | Yes | | | | | | |
| | AIPKLP**S**IDKREA | | | | | | | | Yes |
| | VAGALD**S**TDGAEA | | | | | | | | Yes |
| | AGALDS**T**DGAEAK | | | | | | | Yes | |
| | RASFSS**S**RPLSPR | | | | | | | | Yes |
| | SSSRPL**S**PRTTLP | | Yes | | Yes | | | | |
| *Crassostrea* | PLSRTG**T**PTAADK | | | | Yes | | | | |
| | RAKYVE**T**PVQSCS | | | | | | Yes | | |
| | VETPVQ**S**CSKTRG | | | | | | | | Yes |
| | KLNALK**S**PRKPDG | | Yes | | | | | | |
| *Hemicentrotus* | PLSRTG**S**PTAAEK | | | | Yes | | | | |
| | PINTPA**S**ILSSRN | | | | | | | | Yes |
| | YLRDII**T**GKSEDH | | Yes | | | | | | |

| | | | | | | | | | |
|---|---|---|---|---|---|---|---|---|---|
| | FDNILL**T**MGGKGI | | | | | | | | Yes |
| *Oncorhynchus* | GDVPAP**S**SSPAEK | | | | | | | | Yes |
| | PISRPL**T**PKQQLD | | Yes | | | | | | |
| | DVLGGG**S**GTRRYQ | | | | | | | | Yes |
| | SRGLED**S**QSREAG | | | | | | | Yes | |
| | QPPTCG**S**PTATEK | | | | | | | | Yes |
| *Rattus* | SGIPKA**T**TSAIEK | | | | | | | | Yes |
| Takifugu | EQQQMV**S**PRMRSL | | | | | | Yes | | |
| | TPSRMA**S**VPSEQQ | | | | | | | | Yes |
| | ERRAAM**S**SPPMTP | | | | | | | | Yes |
| *Tetrahymena* | QIDNLK**T**MKQPAE | | Yes | | | | | | |
| *Trypanosoma* | DGKRLA**S**AQGQGD | | | | Yes | | | | |
| ***Inner Arm Group 4 HCs*** | | | | | | | | | |
| *Chlamydomonas* | EPTQEG**S**QASVFD | | | | | | | | Yes |
| | LHQLAH**S**GNGSAE | | | | | | | Yes | |
| | GGSGRQ**S**LTRLAA | | | | | | | | Yes |
| | SGRQSL**T**RLAAYM | | | | | | | | Yes |
| *Crassostrea* | SRMSRS**S**HGGKPF | | | | | | | | Yes |
| | HGGKPF**S**ANRTRV | | | | | | | | Yes |
| *Oncorhynchus* | TTFRTP**S**DLHSLV | | | | | | | | Yes |
| *Rattus* | LGQPRK**S**PLAGTD | | | | | | | | Yes |
| *Trypanosoma* | GQNAPL**S**PGRSEL | | | | | | Yes | | |
| | TATDTV**S**ERPKTA | | Yes | | | | | | |
| ***Inner Arm Group 5 HCs*** | | | | | | | | | |
| *Chlamydomonas* | VRHRYP**S**NNGSAA | | | | Yes | | | | |
| | LDAGRR**S**HPGASA | | | | | | | | Yes |
| | RPTRGP**S**ELGEPA | | | | Yes | | | | |
| | GPVLAR**S**ISQVLP | | | | | | | Yes | |
| | VLARSI**S**QVLPEG | | | | Yes | | | | |
| | SREDQQ**S**PGGLKQ | | | | | | | | Yes |
| | LETLKR**T**PQSPKT | | | | | | | | Yes |
| | LKRTPQ**S**PKTDVP | | Yes | | | | | | |
| | TPQSPK**T**DVPPLD | | | | | | | | Yes |

| | | | | | | | | | |
|---|---|---|---|---|---|---|---|---|---|
| | LDKPLQ**T**APALRG | | | | | | | | Yes |
| | APALRG**S**ARRREG | | Yes | | | | | Yes | |
| | RREGEG**S**PPGALG | | | | | Yes | | | |
| | GLRPED**S**SAEAAQ | | | Yes | | | | | |
| *Crassostrea* | KMRRAG**S**VEPMEK | | | | Yes | | | | |
| | PMEKPE**S**PLPELP | | | | | | | | Yes |
| *Hemicentrotus* | NGSPTR**S**LNLPGV | | | | | | | | Yes |
| | VAADGK**S**SPTGDK | | | | | | | | Yes |
| | GDKDQA**T**TSTNDI | | | | | | | | Yes |
| *Trypanosoma* | LDPLKG**T**LQSSTS | | | | | | | Yes | |
| | LKGTLQ**S**STSGSP | | | | | | | | Yes |
| | RPRGIR**S**PYNPYD | | | | | | | | Yes |
| | GGGGGT**S**PSGVLG | | | | | | | | Yes |
| | MQSKRD**S**SSEFLE | Yes | | Yes | | | | | |
| | LETLYQ**S**TERGKT | | | | | | | Yes | |
| | NKVCDA**S**SLSRER | | | | | | | | Yes |
| ***Unassigned Inner Arm HC*** | | | | | | | | | |
| *Chlamydomonas* | VHGGGG**S**PLRHTG | | | | | | | | Yes |
| | SPGRRV**S**GSISPL | | | | Yes | | | | |
| | GRRVSG**S**ISPLPG | | | | | | | | Yes |
| | RVSGSI**S**PLPGAA | | | | | | | | Yes |
| | AASGAA**S**RGSSPS | | | | | | | | Yes |
| | ATGAGG**S**GDGAAD | | | | | | | | Yes |
| | SLRRPG**S**GGGASG | | | | Yes | | | | |
| | SRGGSR**S**ASPAPG | | | | | | | | Yes |
| | GGSRSA**S**PAPGPP | | | | Yes | | | | |
| | SGGDPA**S**PSPGGI | | | | | | | | Yes |
| | DVIGEG**S**EEALRP | | | | | | | | Yes |
| | TAVASD**S**PRESIF | | Yes | Yes | | | | | |

* Consensus sequences are from ProSite (https://prosite.expasy.org/), Seok (2021) *Life* **11**, 957 and Moniz *et al*. (2011) *Cell Division* **6**, 18. The phosphorylated residues are indicated in **bold**.

#### Table S3 Phosphorylated Sites on *Chlamydomonas* Dynein Heavy Chains Identified in Cytoplasm

| Dynein Heavy Chain | Modified Residue | Sequence Context* | Phospho-Ser/Thr Identified in Cilia |
|---|---|---|---|
| DHC2 (IAD group 4) | $S_{52}$ | EPTQEG**S**QASVFD | Yes |
| DHC4 (IAD group 3) | $S_{197}$ | AIPEEV**S**GASLDY | No |
| | $S_{489}$ | IIARPP**S**PTRTVP | No |
| | $S_{4532}$ | LADEDG**S**EGGGHD | Yes |
| DHC7 (IAD group 5) | $S_{90}$ | LKRTPQ**S**PKTDVP | Yes |
| | $T_{104}$ | LDKPLQ**T**APALRG | Yes |
| | $S_{111}$ | APALRG**S**ARRREG | Yes |
| DHC8 (IAD group 3) | $S_{35}$ | LPGRTG**S**PGAGFT | Yes |
| | $S_{88}$ | AIPKLP**S**IDKREA | Yes |
| DHC9 (IAD group 3) | $S_{159}$ | SSSRPL**S**PRTTLP | Yes |
| DHC11 (IAD group 3) | $S_{142}$ | KHPRPI**S**PSKPRN | No |
| | $S_{250}$ | KLPRIP**S**PGTYAA | No |
| | $S_{1475}$ | HWERLS**S**TLGQRV | No |
| DHC13 (OAD $\alpha$) | $S_{3649}$ | EKARRS**S**GVGDRR | Yes |
| | $S_{3657}$ | VGDRRP**S**QEGLPG | Yes |

* The phosphorylated residue identified is indicated in **Bold**.

**Table S4** **Modification Stoichiometry at Dynein Heavy Chain Phosphorylation Sites Based on Mass Spectral Peak Intensities**

| Organism | Heavy Chain | Phospho-site | Phospho-Peptide Intensity | Unmodified Peptide Intensity | Approx. Amount Modified (%) |
|---|---|---|---|---|---|
| ***Outer Arm Dynein $\alpha$ HC*** | | | | | |
| *Chlamydomonas* | DHC13 | $S_{3630}$ | 3.38E7 | 4.09E7 | 45 |
| | | $S_{3657}$ | 1.12E8 | 1.08E8 | 51 |
| | | $S_{3674}$ | 6.26E7 | 9.53E8 | 6 |
| *Tetrahymena* | DYH5 ($\gamma$ HC) | $S_{3381}$ | 6.10E7 | Not found[&] | 100 |
| ***Outer Arm Dynein $\beta$ HC*** | | | | | |
| *Crassostrea* | DNAH9 | $S_{3}$ | 2.63E7 | 6.21E7 | 30 |
| *Rattus* | DNAH9 | $S_{13}$ | 9.45E6 | 2.04E7 | 32 |
| | DNAH11 | $S_{13}$ | 1.47E7 | 1.67E7 | 47 |
| *Trypanosoma* | Tb927.11.3250 | $S_{1050}$ | 1.33E8 | 3.57E8 | 27 |
| | | $S_{1056}$ | 2.03E7 | 3.57E8 | 6 |
| | | $S_{4089}$ | 1.91E7[$] | Not found | 100 |
| | | $S_{4091}$ | 1.91E7[$] | Not found | 100 |
| ***Outer Arm Dynein $\gamma$ HC*** | | | | | |
| *Ciona* | DNAH8 | $S_{876}$ | 4.64E7 | 1.14E8 | 29 |
| | | $S_{971}$ | 2.39E7 | 6.15E7 | 28 |
| *Crassostrea* | DNAH8 | $S_{29}$ | 2.51E7 | 9.41E6 | 73 |
| | | $S_{48}$ | 2.91E6 | 1.64E7 | 15 |
| *Hemicentrotus* | DNAH8 ($\alpha$ HC) | $T_{1071}$ | 7.63E6 | Not found | 100 |
| | | $S_{1092}$ | 2.73E6 | Not found | 100 |
| ***Inner Arm Dynein I1/f $\alpha$ HC*** | | | | | |
| *Chlamydomonas* | DHC1 | $S_{1050}$ | 8.36E6 | 4.59E8 | 2 |
| *Crassostrea* | DNAH10 | $S_{111}$ | 9.52E6 | 1.00E7 | 49 |
| ***Inner Arm Dynein I1/f $\beta$ HC*** | | | | | |
| *Hemicentrotus* | DNAH2 | $S_{42}$ | 1.59E7 | 8.22E6 | 66 |
| *Takifugu* | DNAH2 | $S_{125}$ | 7.87E6 | 9.66E7 | 7 |
| *Tetrahymena* | DYH7 | $T_{3229}$ | 8.50E7 | 3.06E7 | 74 |

| | | | | | |
|---|---|---|---|---|---|
| ***Inner Arm Group 3 HCs*** | | | | | |
| *Chlamydomonas* | DHC4 | $T_{941}$ | 9.32E6[#] | 4.62E7 | 17 |
| | | $S_{945}$ | 4.98E7[#] | 4.62E7 | 52 |
| | | $S_{4532}$ | 1.24E8 | 5.25E7 | 70 |
| | DHC6 | $T_{128}$ | 3.43E7 | 6.66E7 | 34 |
| | | $T_{205}$ | 3.79E7 | 2.62E8 | 11 |
| | DHC8 | $S_{88}$ | 3.76E8 | Not found | 100 |
| | DHC9 | $S_{155}$ | 3.32E7[+] | 5.47E8 | 6 |
| | | $S_{159}$ | 3.68E8[+] | 5.47E8 | 40 |
| *Crassostrea* | DNAH3 | $T_{11}$ | 1.16E7 | 2.89E6+6.59E6[#] | 54 |
| | | $S_{15}$ | 6.59E6 | 2.89E6+1.16E7[#] | 45 |
| | | $S_{107}$ | 1.2E7 | Not found | 100 |
| | DNAH7 | $T_{142}$ | 9.75E7 | 9.91E7 | 50 |
| *Hemicentrotus* | DNAH7 | $S_{147}$ | 3.47E7 | 3.42E7 | 50 |
| | DNAH12 | $S_{57}$ | 4.28E6 | 8.42E6 | 34 |
| | | $T_{1593}$ | 5.15E7 | Not found | 100 |
| | | $T_{3405}$ | 2.11E6 | 1.07E8 | 2 |
| *Oncorhynchus* | DNAH3 | $S_{126}$ | 8.65E6 | 1.12E7 | 7 |
| | | $T_{169}$ | 1.63E7 | Not found | 100 |
| | DNAH7 | $S_{70}$ | 1.73E6 | Not found | 100 |
| | | $S_{85}$ | 1.18E7 | 2.48E7 | 32 |
| | | $S_{144}$ | 2.03E7 | 3.53E6 | 85 |
| *Rattus* | DNAH7 | $T_{152}$ | 5.76E6 | 6.60E7 | 8 |
| *Takifugu* | DNAH3 | $S_{68}$ | 3.65E7 | 6.39E8 | 5 |
| | | $S_{78}$ | 6.40E7 | 6.39E8 | 9 |
| | | $S_{120}$ | 1.03E8 | 1.79E8 | 36 |
| *Tetrahymena* | DYH25 | $T_{1092}$ | 9.46E7 | 3.21E6 | 97 |
| *Trypanosoma* | Tb927.2.5270 | $S_{152}$ | 1.09E7 | 3.06E7 | 26 |
| ***Inner Arm Group 4 HCs*** | | | | | |
| *Chlamydomonas* | DHC2 | $S_{52}$ | 1.32E7 | 1.41E8 | 8 |
| | | $S_{192}$ | 2.10E7 | 2.47E8 | 8 |
| | | $S_{2480}$ | 7.91E7[$] | Not found | 100 |
| | | $T_{2482}$ | 7.91E7[$] | Not found | 100 |

| | | | | | |
|---|---|---|---|---|---|
| *Crassostrea* | DNAH1 | $S_{9}$ | 2.44E6[$] | Not found | 100 |
| | | $S_{16}$ | 2.44E6[$] | Not found | 100 |
| *Oncorhynchus* | DNAH1 | $S_{100}$ | 7.24E6 | 7.44E7 | 49 |
| *Rattus* | DNAH1 | $S_{65}$ | 4.31E7 | 1.32E7 | 77 |
| *Trypanosoma* | Tb11.10.5350 | $S_{26}$ | 1.18E7 | 4.70E7 | 20 |
| | Tb927.11.8160 | $S_{49}$ | 1.87E7 | 3.39E8 | 5 |
| ***Inner Arm Group 5 HCs*** | | | | | |
| *Chlamydomonas* | DHC3 | $S_{152}$ | 8.51E6 | Not found | 100 |
| | | $S_{171}$ | 3.78E7 | Not found | 100 |
| | | $S_{685}$ | 4.62E7 | 1.27E8 | 27 |
| | | $S_{5326}$ | 3.18E8 | Not found | 100 |
| | DHC7 | $S_{9}$ | 1.07E7 | 7.01E8 | 1 |
| | | $S_{90}$ | 2.32E8 | Not found | 100 |
| | | $T_{104}$ | 3.12E7 | 1.86E7 | 63 |
| | | $S_{111}$ | 5.40E8 | 6.08E8 | 47 |
| *Crassostrea* | DNAH6 | $S_{137}$ | 9.46E6+2.79E7[@] | 2.90E7+2.79E7 | 40 |
| | | $S_{146}$ | 2.34E7+2.79E7[@] | 2.90E7+9.46E6 | 57 |
| *Hemicentrotus* | DNAH6 | $S_{19}$ | 8.39E6 | 1.35E8 | 6 |
| | | $S_{1390}$ | 1.07E7* | 8.01E6* | 57 |
| | | $T_{1400}$ | 8.01E6* | 1.07E7* | 43 |
| *Trypanosoma* | Tb927.11.11220 | $T_{155}$ | 5.35E6 | 2.35E7 | 18 |
| | | $S_{158}$ | 1.91E7 | 5.67E7 | 25 |
| | | $S_{222}$ | 1.78E7 | 2.40E8 | 7 |
| | | $S_{3173}$ | 2.98E6 | 3.02E8 | 1 |
| | Tb927.7.920 | $S_{7}$ | 1.05E7 | 6.82E7 | 15 |
| | | $S_{18}$ | 7.23E6 | 7.66E7 | 9 |
| | | $S_{75}$ | 6.82E6 | 7.20E6 | 49 |
| ***Unassigned Inner Arm HC*** | | | | | |
| *Chlamydomonas* | DHC12 | $S_{758}$ | 1.57E7 | 4.13E7 | 28 |
| | | $S_{1124}$ | 1.13E7 | 1.06E7 | 52 |
| | | $S_{4848}$ | 8.76E6 | 1.35E7 | 39 |
| ***Cytoplasmic Dynein HC*** | | | | | |
| *Rattus* | DYNC1H1 | $S_{634}$ | 4.70E6 | Not found | 100 |

& In cases where the unmodified peptide was not found, the amount of the phosphorylated form was set to 100%.

* These derive from long peptides that had either pSer or pThr. The fully unmodified form was not found.

### These sites occur in the same peptide. Both mono-phosphorylated forms and the unmodified peptide were found.

$ These sites occur in the same peptide. Only the doubly phosphorylated form was identified.

@ These sites occur in the same peptide. Double-, both single- and un-phosphorylated peptides were found.

+ These sites occur in the same peptide. The unmodified, one mono-phosphorylated and the doubly phosphorylated forms were found.

**Table S5** **Modification Stoichiometry at Outer Arm Docking Complex Phosphorylation Sites in Wildtype (CC-125) and *uni1-1* Mutant Cilia Based on Mass Spectral Peak Intensities**

| Docking Complex Component | Phospho-site | Phospho-Peptide Intensity | Unmodified Peptide Intensity | Approx. Amount Modified (%) |
|---|---|---|---|---|
| **DC1 (CC-125)** | | | | |
| | $S_{33}$ | 1.02E7 | 3.72E7 | 22 |
| | $T_{73}$ | 2.31E7 | 1.47E8 | 14 |
| | $T_{351}$ | 1.42E7 | 4.67E8 | 3 |
| | $T_{498}$ | 3.85E7 | 1.22E8 | 24 |
| | $T_{508}$ | 3.22E7 | 5.57E8 | 5 |
| | $S_{510}$ | 6.31E7 | 5.57E8 | 10 |
| | $S_{512}$ | 2.16E7*[1] | 5.57E8 | 4 |
| | $S_{514}$ | 2.16E7*[1] | 5.57E8 | 4 |
| | $S_{521}$ | 2.16E7*[1] | 5.57E8 | 4 |
| | $S_{628}$ | 3.66E7 | 7.80E8 | 4 |
| | $T_{698}$ | 2.10E8 | Not found[&] | 100 |
| | $S_{700}$ | 3.14E7*[2] | Not found | 100 |
| | $S_{709}$ | 3.14E7*[2] | Not found | 100 |
| **DC2 (CC-125)** | | | | |
| | $S_{12}$[#] | 1.23E8*[1] | 9.30E7 | 57 |
| | $S_{15}$ | 6.09E7*[1] | 9.30E7 | 40 |
| | $T_{20}$ | 1.23E8*[1] | 9.30E7 | 57 |
| | $S_{266}$ | 2.44E7*[2] | 1.16E8 | 17 |
| | $S_{267}$ | 2.19E8*[2] | 1.16E8 | 65 |
| | $T_{512}$ | 3.89E7 | 6.59E8 | 56 |

| | | | | |
|---|---|---|---|---|
| | $S_{548}$ | 7.54E6 | 6.92E8 | 1 |
| **DC3 (CC-125)** | | | | |
| | No sites found | | | |
| **DC1 (*uni1-1*)** | | | | |
| | $S_{347}$ | 3.88E6 | 6.87E7 | 5 |
| | $T_{652}$ | 1.05E7 | 2.07E7 | 33 |
| **DC2 (*uni1-1*)** | | | | |
| | $S_{12}$ | 2.01E7*[1] | Not found | 100 |
| | $S_{15}$ | 2.01E7*[1] | Not found | 100 |
| | $S_{266}$ | 1.02E7 | 1.74E7 | 36 |
| **DC3 (*uni1-1*)** | | | | |
| | No sites found | | | |

& In cases where the unmodified peptide was not found, the amount of the phosphorylated form was set as 100%.

*n Indicates clusters present on the same peptide.

### The N-terminal sequence for DC2 shown in the current version (CC-4532 v.6.1) of the *Chlamydomonas* genome is incorrect. Our previous mass spectral analysis (Sakato-Antoku *et al*. *Cells* **12**, 2492 [2023]) identified the N-terminal sequence processed by methionine aminopeptidase as (M)PSADATR that was indicated in the previous genome version (v.5.6). Thus, the residue numbering used here reflects the experimentally confirmed DC2 sequence from genome v.5.6; both v.5.6 and v.6.1 genomes are available at Phytozome.
